# Alignment of auditory artificial networks with massive individual fMRI brain data leads to generalizable improvements in brain encoding and downstream tasks

**DOI:** 10.1101/2023.09.06.556533

**Authors:** Maelle Freteault, Maximilien Le Clei, Loic Tetrel, Pierre Bellec, Nicolas Farrugia

## Abstract

Artificial neural networks trained in the field of artificial intelligence (AI) have emerged as key tools to model brain processes, sparking the idea of aligning network representations with brain dynamics to enhance performance on AI tasks. While this concept has gained support in the visual domain, we investigate here the feasibility of creating auditory artificial neural models directly aligned with individual brain activity. This objective raises major computational challenges, as models have to be trained directly with brain data, which is typically collected at a much smaller scale than data used to train AI models.

We aimed to answer two key questions: (1) Can brain alignment of auditory models lead to improved brain encoding for novel, previously unseen stimuli? (2) Can brain alignment lead to generalisable representations of auditory signals that are useful for solving a variety of complex auditory tasks? To answer these questions, we relied on two massive datasets: a deep phenotyping dataset from the Courtois neuronal modelling project, where six subjects watched four seasons (36 hours) of the *Friends* TV series in functional magnetic resonance imaging and the HEAR benchmark, a large battery of downstream auditory tasks.

We fine-tuned SoundNet, a small pretrained convolutional neural network with ∼2.5M parameters. Aligning SoundNet with brain data from three seasons of *Friends* led to substantial improvement in brain encoding in the fourth season, extending beyond auditory and visual cortices. We also observed consistent performance gains on the HEAR benchmark, particularly for tasks with limited training data, where brain-aligned models performed comparably to the best-performing models regardless of size. We finally compared individual and group models, finding that individual models often matched or outperformed group models in both brain encoding and downstream task performance, highlighting the data efficiency of fine-tuning with individual brain data.

Our results demonstrate the feasibility of aligning artificial neural network representations with individual brain activity during auditory processing, and suggest that this alignment is particularly beneficial for tasks with limited training data. Future research is needed to establish whether larger models can achieve even better performance and whether the observed gains extend to other tasks, particularly in the context of few shot learning.

## Introduction

### Overall objective

Artificial neural networks (ANNs) have emerged as a powerful tool in cognitive neuroscience. Specifically, ANNs trained to solve complex tasks directly from rich data streams, such as natural images, are able to accurately encode brain activity, i.e. predict brain responses directly from the stimulus. A notable observation is that the ANNs which have high performance solving behavioural tasks, e.g. image classification, tend to be the ones which perform best to predict brain activity. This was noted first in vision (Yamins et al., 2014) and rigorously established for language while controlling for model architecture and model capacity (Caucheteux et al., 2023). This result suggests that directly training the representations of ANNs to encode well brain activity may lead to more generalizable representations and improved performance on novel downstream tasks. This process, in general called brain alignment (Sucholutsky et a., 2023, Mineault et al., 2024), has only been explored in a few works so far, and most of these works have been carried in the field of vision (Seeliger et al., 2021; St-Yves et al., 2023; Lu et al., 2024) and language (Konkle et al., 2022; Schwartz et al., 2019). These brain alignment works used datasets of limited size, both for brain encoding and downstream tasks. In this work, we explore for the first time the impact of individual brain alignment on an auditory ANN. The study also leverages massive datasets both for training the networks, testing the generalisation of brain encoding to novel stimuli, with the Courtois NeuroMod fMRI dataset (Boyle et al., 2020), and evaluating task performance on a wide range of downstream tasks, with the HEAR benchmark (Turian et al., 2022).

### ANNs in audio classification and brain encoding

ANNs trained with deep learning for cognitive neuroscience emerged initially in the field of vision (Schrimpf et al., 2020). CNNs in artificial vision share strong parallels with visual brain processing (Bengio et al., 2013; Krizhevsky et al., 2012). It was found that ANNs trained on complex tasks such as image annotation could predict brain responses to image stimuli with considerable accuracy (Yamins et al., 2014). Building on these foundations, the capability of ANNs for brain encoding has been extended to both language and auditory neuroscience. Kell and collaborators designed an auditory CNN with two branches, tailored for music and speech recognition (Kell et al., 2018). They discovered distinct auditory processing streams in their network, with the primary auditory cortex best predicted by middle layers, and the secondary auditory cortex by late layers. More recently, Giordano and coll. provided further evidence for the intermediary role of the superior temporal gyrus (STG) in auditory processing using 3 different CNNs (Giordano et al., 2023), while Caucheteux and coll. use of a modified GPT-2 provided new insights into predictive coding theory (Caucheteux et al., 2023).

### Brain alignment

Typical brain encoding studies re-use a pretrained network based on a very large collection of sounds, and can feature a very large number of parameters. It is relatively straightforward to apply such large networks for brain encoding, e.g. using Ridge regression from the latent space of the network to brain activity, as it was done for example with BERT (Devlin et al., 2019). Aligning internal representations of ANNs models with brain activity is a much more ambitious goal, which requires directly optimising the parameters of a network in order to maximise the quality of brain encoding through a process called fine-tuning. This optimization process raises a number of computational and conceptual challenges. First, there is clear evidence of substantial inter-individual differences in functional brain organisation (Gordon et al., 2017; Gratton et al., 2018). For this reason, some authors have advocated for building individual brain models using deep fMRI datasets, where a limited number of individuals get scanned for an extended time, instead of datasets featuring many subjects with limited amounts of data per subject (Naselaris et al., 2021). It is the approach which we decided to take. Second, most fMRI datasets feature only a limited number of stimuli which can be used to train a network. The largest fMRI stimuli sets include Dr Who (approximately 23 hours of video stimuli) (Seeliger et al., 2019) and Natural scenes dataset (10k images) (Allen et al., 2022), which is orders of magnitude smaller than what is currently used in the AI field. For example, the latest version of the recent auditory ANN wav2vec (Baevski et al., 2020) has 317 millions parameters, and was pretrained with over 60k hours of sound data. It thus seems likely that network architectures should feature less parameters when trained for brain alignment than state-of-art networks trained in the field of Artificial Intelligence (AI), and the few published studies of brain alignment indeed followed this trend, e.g. (Seeliger et al., 2021; St-Yves et al., 2023).

### Brain alignment and generalisation of behaviour

In this work, the term “brain alignment” refers to optimizing the parameters of a pretrained ANN to improve brain encoding performance, as outlined in the previous paragraph. In contrast, the term “human alignment*”*, commonly used in the field of AI safety, refers to ensuring that artificial systems behave in accordance with human intentions, expectations and benefits (Ji et al., 2023). A recent white paper by Mineault et al. (2024) highlights the potential of brain alignment to enhance human alignment in the context of AI safety, through increased robustness of behaviour. This avenue was further discussed at a recent workshop of the 2024 international conference on representational learning (ICLR) (Sucholutsky et a., 2023). However, given the limited size of neuroimaging datasets, compared to datasets commonly used to train AI models, it is not clear that fine-tuning can lead to generalizable performance gains, and may on the contrary distort pre-trained features to overfit training data with poor out-of-distribution performance (Geirhos et al., 2020, Kumar et al., 2022). A few previous studies still found that brain alignment may improve the behaviour of ANNs on downstream tasks (Palazzo et al., 2020; Nishida et al., 2020, Moussa et al., 2024)— tasks with available ground truth to assess performance and that the ANN was not explicitly trained to perform. However, these works examined only a limited number of downstream tasks (one in Palazzo et al. and four in Nishida and Moussa) and reported, at best, modest performance gains. These limitations may be attributed to the small size of the datasets used for brain alignment or the narrow scope of the downstream tasks considered.

### Generalisation scope of brain encoding models

While brain encoding studies are rapidly gaining traction in computational cognitive neuroscience, the scope of generalisation tested in these models remains limited. For instance, seminal work on auditory brain encoding used only 165 sounds, each 2 seconds long (Kell et al., 2018), and pioneering studies on language comprehension often relied on less than 30 minutes of recordings per subject, typically featuring a single story (Caucheteux et al., 2023). In vision, the Natural Scene Dataset (Allen et al., 2022) included ∼10,000 image stimuli per subject for a visual recognition task; but these images spanned only about 40 semantic categories (Shirakawa et al., 2024). In this study, we aimed to broaden the scale of brain encoding generalisation by using the audio tracks of three full seasons of the *Friends* TV show for training (28 hours), and an additional, separate season for testing (9 hours). These complex audio stimuli included extensive speech from a large and diverse set of speakers, interwoven with music and a variety of naturalistic sounds. Additionally, we aimed to investigate a broad range of downstream behavioural tasks to assess how brain alignment using *Friends* stimuli influences different aspects of sound processing.

### Courtois NeuroMod, HEAR benchmark and model architecture

In this work, we aimed at demonstrating the feasibility of brain alignment of artificial neural networks in the auditory domain. Specifically, we addressed two general questions: (1) Do brain-aligned networks encode brain activity related to auditory processing more effectively when using novel stimuli? (2) Do brain-aligned networks show improved performance in downstream auditory tasks that are unrelated to the stimuli used for brain encoding?

We made several key design choices to address these two questions:

First, following the approach advocated by Naselaris et al. (2021), we decided to align ANNs at the individual level and compared their performance to ANNs trained at the group level. This allowed us to evaluate whether a smaller but individual-specific dataset is more advantageous than a larger group dataset. To achieve this, we leveraged the Courtois NeuroMod dataset (Boyle et al., 2020), the largest deep fMRI dataset to date, which was specifically designed by our team to align ANNs with brain data. Its 2022 release features a small number of subjects (N=6), each with over 100 hours of fMRI data, with additional data yet to be released. The dataset spans a wide range of tasks across multiple domains, including several movie-watching datasets with complex soundtracks. For this study, we focused on the *Friends* dataset, which is both extensive (36 hours of data per subject) and varied (covering multiple seasons of the TV show with different stimuli in each episode).

Second, in terms of model size and architecture, we selected a pretrained model called SoundNet, which has been shown to perform well in sound processing and brain encoding (Farrugia et al., 2019; Nishida et al., 2020). Soundnet features a limited number of parameters (fewer than 3 millions) as well as a simple convolutional architecture with decreasing temporal resolution, making it well-suited to fine-tuning with an fMRI dataset. The *Friends* dataset allowed us to test the generalisation of brain-aligned models to new stimuli in a large controlled distribution (a different season of *Friends*) and to replicate the process of brain alignment with six different subjects.

Third, to assess generalisation abilities on downstream tasks, we leveraged a recent machine learning competition : the Holistic Evaluation of Auditory Representations (HEAR, Turian et al., 2022). HEAR offers a standardised procedure to test the generalisation of the internal representations of a model on a wide array of downstream tasks. Using the HEAR environment, we evaluated our brain-aligned models and, given the large number of teams that participated in the HEAR competition, were able to rigorously compare their performance against a range of state-of-the-art AI approaches.

### Study objectives

The specific objectives and hypotheses of our study are as follows:

- Align SoundNet with individual and group brain data and compare the quality of brain encoding with the baseline, non-brain-aligned model. Our hypothesis was that the alignment procedure would lead to substantial gains in brain encoding for within-distribution stimuli drawn from the *Friends* dataset, and that these gains would be subject-specific, i.e. they would not transfer to other individuals.
- Evaluate how brain alignment impacts out-of-distribution downstream tasks. Our hypothesis was that brain alignment would lead to no degradation or even modest improvements in performance across a wide range of tasks.

Taken together, this study establishes the feasibility and some key methodological decisions to align auditory networks with brain data using massive individual fMRI data, and clarifies how this alignment impacts the performance of networks on downstream tasks.

## Materials and Methods

### fMRI data

#### Participants

Six healthy participants (aged 31 to 47 at the time of recruitment in 2018), 3 women (sub-03, sub-04 and sub-06) and 3 men (sub-01, sub-02 and sub-05) were recruited to participate in the Courtois Neuromod Project for at least 5 years. All subjects provided informed consent to participate in this study, which was approved by the ethics review board of the “CIUSS du centre-sud-de-l’île-de-Montréal” (under number CER VN 18-19-22). Three of the participants reported being native francophone speakers (sub-01, sub-02 and sub-04), one as being a native anglophone (sub-06) and two as bilingual native speakers (sub-03 and sub-05). All participants reported the right hand as being their dominant hand and reported being in good general health. Exclusion criteria included visual or auditory impairments that would prevent participants from seeing and/or hearing stimuli in the scanner and major psychiatric or neurological problems. Standard exclusion criteria for MRI and MEG were also applied. Lastly, given that all stimuli and instructions are presented in English, all participants had to report having an advanced comprehension of the English language for inclusion. The above boilerplate text is taken from the cNeuroMod documentation^1^ (Boyle et al., 2020), with the express intention that users should copy and paste this text into their manuscripts unchanged. It was released by the Courtois NeuroMod team under the CC0 license.

#### *Friends* Dataset

The dataset used in this study is a subset of the 2022-alpha release of the Courtois Neuromod Dataset, called *Friends*, where the participants have been watching the entirety of seasons 1 to 4 of the TV show *Friends*. This subset was selected because it provided a rich naturalistic soundtrack, with both a massive quantity of stimuli and a relative homogeneity in the nature of these stimuli, as the main characters of the series remain the same throughout all seasons. Subjects watched each episode cut in two segments (a/b), also referred as *runs*, to allow more flexible scanning and give participants opportunities for breaks. There is a small overlap between the segments to allow participants to catch up with the storyline. The episodes were retransmitted using an Epson Powerlite L615U projector that casted the video through a waveguide onto a blank screen located in the MRI room, visible to the participants via mirror attached to the head coil. Participants wore MRI compatible S15 Sensimetric headphone inserts, providing high-quality acoustic stimulation and substantial attenuation of background noise, and wore custom sound protection gear. More details can be found on the Courtois Neuromod project website^1^.

#### Data acquisition

The participants have been scanned using a Siemens Prisma Fit 3 Tesla, equipped with a 2-channel transmit body coil and a 64-channel receive head/neck coil. Functional MRI data was acquired using an accelerated simultaneous multi-slice, gradient echo-planar imaging sequence (Xu et al., 2013) : slice acceleration factor = 4, TR = 1.49 s, TE = 37 ms, flip angle = 52 degrees, voxel size = 2 mm isotropic, 60 slices, acquisition matrix 96×96. In each session, a short acquisition (3 volumes) with reversed phase encoding direction was run to allow retrospective correction of B0 field inhomogeneity-induced distortion.

To minimise head movement, the participants have been provided individual head cases adapted to the shape of their head. Most imaging in the Courtois Neuromod project are composed solely of functional MRI runs. Periodically, an entire session is dedicated to anatomical scans, see details on the Courtois Neuromod project website^1^. Two anatomical sessions were used per subject in this study for fMRIprep anatomical alignment, specifically a T1-weighted MPRAGE 3D sagittal sequence (duration 6:38 min, TR = 2.4 s, TE = 2.2 ms, flip angle = 8 deg, voxel size = 0.8 mm isotropic, R=2 acceleration) and a T2-weighted FSE (SPACE) 3D sagittal sequence (duration 5:57 min, TR = 3.2 s, TE = 563 ms, voxel size = 0.8 mm isotropic, R=2 acceleration).

#### Preprocessing of the fMRI data

All fMRI data from the 2022-alpha release were preprocessed using the fMRIprep pipeline version 20.2.5 (“long-term support”) (Esteban et al., 2019), see Annex A for details. We used a volume-based spatial normalisation to standard space (MNI152NLin2009cAsym). The anatomical mask derived from the data preprocessing phase was used to identify and select brain voxels. Voxel-level 2D data matrices (TR x voxels) were generated from 4-dimensional fMRI volumes using the NiftiMasker tool from Nilearn (Abraham et al., 2014) and a mask of the bilateral superior temporal gyri middle (middle STG), specifically parcel 153 and 154 of the MIST parcellation (Urchs et al., 2019), resulting in 556 voxels. ROI-level 2D data matrices (TR x ROI) were generated from 4-dimensional fMRI volumes using the NiftiLabelsMasker tool from Nilearn with the MIST parcellation. The MIST parcellation was used as a hard volumetric functional parcellation because of the availability of anatomical labels for each parcel. This functional brain parcellation was also found to have excellent performance in several ML benchmarks on either functional or structural brain imaging (Dadi et al., 2020; Hahn et al., 2022; Mellema et al., 2022). We chose the 210 resolution of the parcellation atlas because parcels were enforced to be spatially contiguous, and separate regions in the left and right hemisphere. Both the middle STG mask used to select the voxels and the parcels from MIST were based on non-linear alignment. For the voxel-level data matrices, we choose to investigate the effect of spatial smoothing by using BOLD time series with no spatial smoothing or smoothed spatially with a 5mm Gaussian kernel. For ROI-level data matrices, we only used BOLD time series with spatial smoothing (5mm gaussian kernel). A so-called “Minimal” denoising strategy was used to remove confounds without compromising the temporal degrees of freedom, by regressing out basic motion parameters, mean white matter, and cerebrospinal fluid signals as available in the library load_confounds^2^ (equivalent to the default denoising strategy now implemented with load_confounds in Nilearn). This strategy is recommended for data with low levels of motion, as is the case for the Courtois NeuroMod sample (Wang et al., 2023).

### Encoding models

#### Overview

Our approach to training encoding models of auditory activity relied on transfer learning and the fine-tuning of a pretrained deep learning backbone, SoundNet. Audio waveforms served as inputs to the backbone, producing as an output a set of time-dependent features. These features were then used to train a downstream convolutional “encoding” layer to predict the fMRI signal. We explored several training variants, including a baseline where only the encoding layer was trained, as well as fine-tuning experiments where SoundNet parameters were updated up to a certain depth in the network, starting with the final layer (see Figure 2). Details of the backbone, fMRI encoding layer, hyperparameters, and training procedures are provided below. Models were implemented using PyTorch and other Python^3^ libraries and were trained on the Alliance Canada infrastructure with V100 and A100 GPUs.

**Figure 1.**
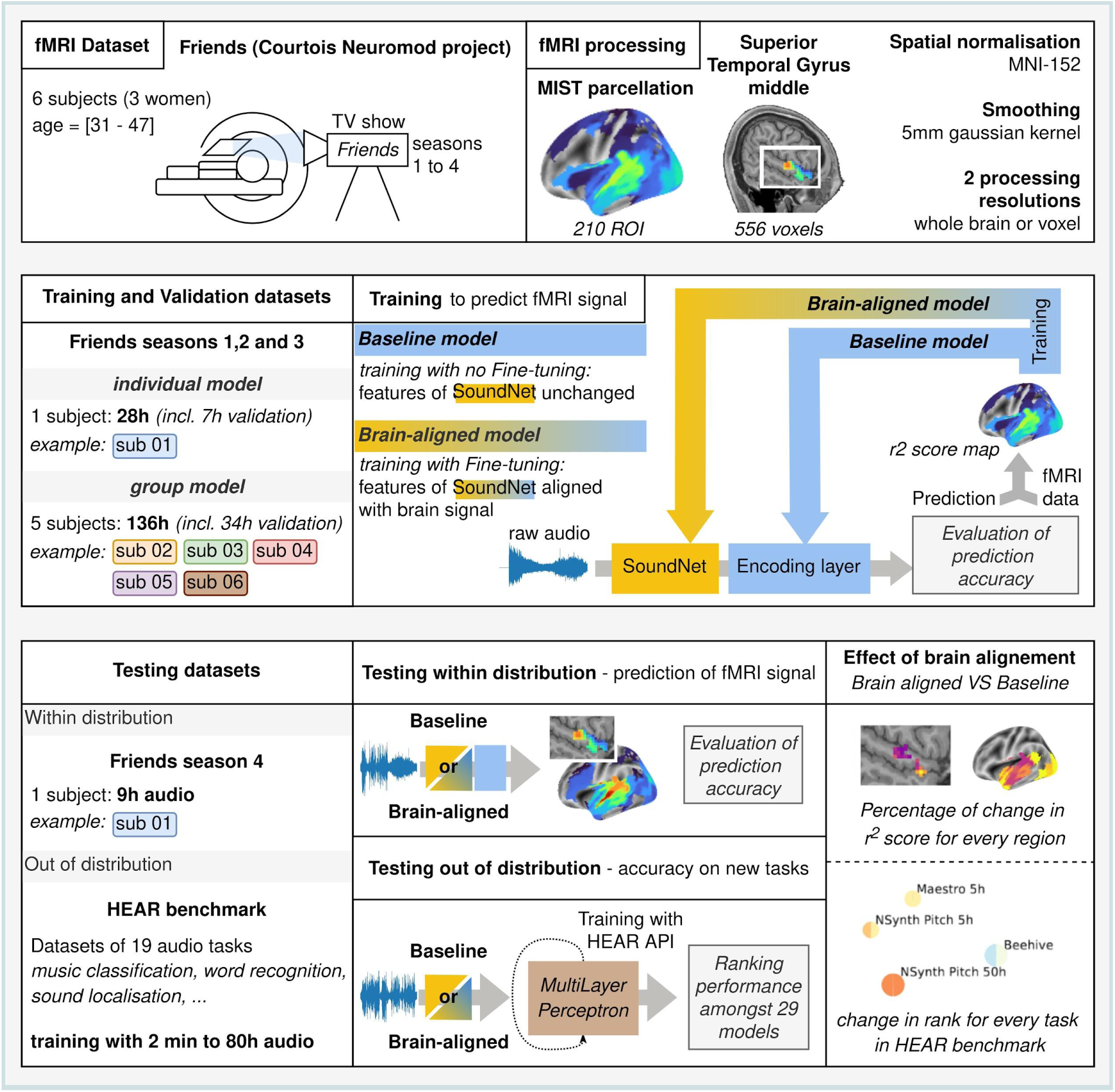
Overview of the analysis. In this study, we used a naturalistic fMRI dataset to align internal features of a pretrained network with brain signals, using an AI training technique called fine-tuning. We evaluated how brain alignment changed the performance of the network, both for tasks that the network has been trained and optimised for (within distribution) and for new tasks (out of distribution).

**Figure 2.**
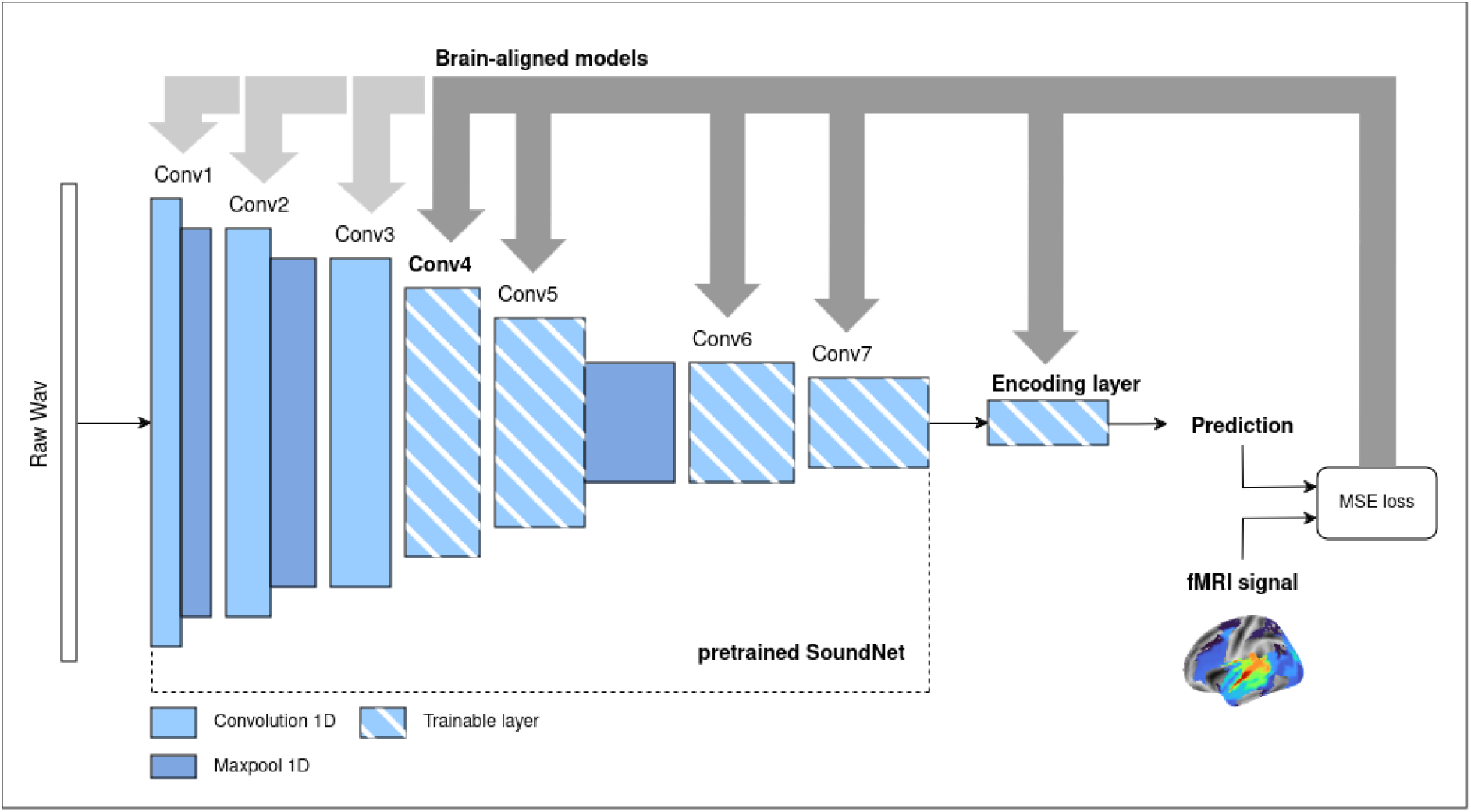
Overview of the training framework. We provided the audio track of the TV show *Friends* to a pretrained convolutional network, SoundNet. Initially, we extracted the output from the 7th convolutional layer of SoundNet, with its parameters *frozen* (fixed values that remain unchanged during training), and used this output as input to train a final encoding layer to predict fMRI activity from a subject watching the TV show. This model serves as our *baseline*. In a second phase, we partially retrained SoundNet along with the encoding layer by fine-tuning all parameters up to the selected layer, allowing these parameters to be updated during training. This new model, where internal layers are fine-tuned to better align with cerebral activity, is referred to as the *brain-aligned* model. The results presented here were obtained using the model fine-tuned up to convolutional layer 4, as depicted in this figure, but we also tested models fine-tuned at various depths, ranging from Conv7 to Conv1.

#### Deep learning backbone

The network we selected as our backbone is SoundNet, a convolutional neural network with the goal of identifying audio content in an audio excerpt, proposed by Aytar et al. (2016). SoundNet was trained to combine information from both the audio and visual inputs of a video, by minimising the Kullback-Leibler divergence (Kullback & Leibler, 1951) between the distribution of its own outputs derived purely from the audio signal, and the output distribution of 2 different vision networks, obtained with the frames of the video. We selected SoundNet for the following reasons : (1) SoundNet is fully convolutional, as all intermediate representations (i.e. layers) are obtained from 1D convolutions and pooling operators, using directly the audio waveform as input with no additional operations. (2) SoundNet was initially trained on a large dataset of natural videos from the Internet (Flickr), with a high degree of correspondence between the visual and audio content, and (3) SoundNet obtained good performances on downstream auditory tasks using transfer learning, as well as good performance as a brain encoding model (Farrugia et al., 2019, Nishida et al., 2020).

At the time of its release in 2016, SoundNet achieved similar performances to the State-of-the-Art (SotA) networks on audio classification benchmarks *Detection and Classification of Acoustic Scenes and Events* DCASE (Mesaros et al., 2017) and *Environmental Sound Classification* ESC-50 (Piczak, 2015; Arandjelovic & Zisserman, 2017). With numerous innovations happening in the AI research field since 2016, as well as the introduction of the dataset AudioSet (Gemmeke et al., 2017), it has since been surpassed by other networks. However, the CNN architecture dominated the leaderboard of many benchmarks up until 2021 for the audio classification task (Wang et al., 2021; Verbitskiy et al., 2022; Gong et al., 2021), and is still quite relevant with the current considerations of the field to find efficient architectures with fewer parameters and reduced energy consumption (Schmid, Koutini, & Widmer, 2023).

While the naturalistic fMRI dataset we used to fine-tune the network is the biggest up-to-date in the fMRI field, the size of the training dataset is still far below what has been offered by larger audio datasets such as AudioSet or VGGSound (Gemmeke et al., 2017; Chen et al., 2020), often used to train the most recent SotA networks. For this reason, we consider that a smaller, simple network was not only necessary in the context of this study, but beneficial to isolate the impact of brain alignment on network performance. As such, SoundNet provides a generic convolutional network to learn from, with its representations encoding audio features of varying durations, and increasing abstraction in deeper layers.

SoundNet’s architecture (Figure 2 and Table 1) is a series of convolutional blocks that always include the following steps :

- a 1D Convolutional layer
- a 1D Batch normalisation realised on the output of the convolutional layer
- a rectified linear unit function element-wise ReLU.

**Table 1.** Architecture of the SoundNet network with the two different encoding layers. The values for the number of parameters shown on the right side of the table have been estimated for Sub-03 encoding models, with a selected duration of 70 TRs and a kernel size of 5 for both encoding layers. Values vary slightly for each subject (see Annex Table 1 for the different values used for each subject).

| Initial training input size<br>Audio Input length * TR length (1,49s) * 22050 Hz |  |  |  |  |  | For Sub-03 :<br>Audio Input length = 70 TR |
| --- | --- | --- | --- | --- | --- | --- |
| Layers |  | Number of out<br>Channels | Kernel size | Stride | Padding | Number of parameters |
| **SoundNet** |  |  |  |  |  |  |
| **Conv1** | Conv1D | 16 | 64 | 2 | 32 | 1040 |
|  | BatchNorm1D | 16 |  |  |  | 32 |
|  | ReLU | 16 |  |  |  |  |
|  | MaxPool1D | 16 | 8 | 8 |  |  |
| **Conv2** | Conv1D | 32 | 32 | 2 | 16 | 16416 |
|  | BatchNorm1D | 32 |  |  |  | 64 |
|  | ReLU | 32 |  |  |  |  |
|  | MaxPool1D | 32 | 8 | 8 |  |  |
| **Conv3** | Conv1D | 64 | 16 | 2 | 8 | 32,832 |
|  | BatchNorm1D | 64 |  |  |  | 128 |
|  | ReLU | 64 |  |  |  |  |
| **Conv4** | Conv1D | 128 | 8 | 2 | 4 | 65,664 |
|  | BatchNorm1D | 128 |  |  |  | 256 |
|  | ReLU | 128 |  |  |  |  |
| **Conv5** | Conv1D | 256 | 4 | 2 | 2 | 131,328 |
|  | BatchNorm1D | 256 |  |  |  | 512 |
|  | ReLU | 256 |  |  |  |  |
|  | MaxPool1D | 256 | 4 | 4 |  |  |
| **Conv6** | Conv1D | 512 | 4 | 2 | 2 | 524,800 |
|  | BatchNorm1D | 512 |  |  |  | 1,024 |
|  | ReLU | 512 |  |  |  |  |
| **Conv7** | Conv1D | 1024 | 4 | 2 | 2 | 2,098,176 |
|  | BatchNorm1D | 1024 |  |  |  | 2,048 |
|  | ReLU | 1024 |  |  |  |  |
| **Encoding layer Whole-brain** |  |  |  |  |  |  |
|  | Conv1D | 210 | Kernel size | 1 | Kernel size -1 | 1,075,410 |
| **Encoding layer STG** |  |  |  |  |  |  |
|  | Conv1D | 556 | Kernel size | 1 | Kernel size -1 | 2,847,276 |

In some of the blocks, a 1D max pooling is also applied to the output of the preceding steps (see Table 1 for more details).

#### fMRI encoding layer

We implemented the SoundNet architecture followed by the fMRI encoding layer as a fully end-to-end trainable network (see Table 1), adapting an open-source PyTorch implementation of SoundNet^4^. Our encoding model predicts entire segments of fMRI data based on corresponding segments of audio waveform data. The SoundNet model is convolutional and non-causal, meaning the output of a filter at a given time point can depend on future time points in the original time series. Accordingly, we designed our encoding layer as a traditional non-causal temporal 1D convolutional layer, with a separate temporal kernel learned for each pair of input features from SoundNet and output features of the brain (either parcel or voxel). The outputs of each feature map after convolution are summed to predict brain activity for a specific brain parcel or voxel. For brain encoding, we used the output of SoundNet’s Conv7 layer, as it matches the temporal resolution of the fMRI signal (0.67 Hz). Formally, the brain encoding layer applies the following model:

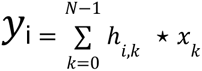

where:

- *y_i_* is a window of brain activity associated with parcel/voxel *i*,
- *N* = 2048 is the number of features in layer 7 (and *k* denoting a particular feature),
- *h_i,k_* is a convolution kernel, with varying size and parameters trainable specifically for each pair (*i*, *k*),
- *x_k_* is a temporal window of activity for the *k*-th feature of layer 7 of SoundNet, and
- * is the valid 1D cross-correlation operator (including padding with a size relative to the kernel size, varying between 6 to 9 seconds, or 4 to 6 TR).

Note that we explored multiple lengths (from 20 s up to 130 s) for the temporal window which we treated as hyper-parameters for optimization (see Annex B on Hyper-parameters exploration).

Notably, with a kernel size of 1, this model is equivalent to a traditional mass-univariate regression of SoundNet features onto brain activity, with no delay between sound waves and brain responses. While the proposed model does not explicitly incorporate an HRF, using a 1D convolution operator with a kernel larger than 1 is analogous to modelling the HRF with a Finite Impulse Response (FIR) filter (Goutte, Nielsen, & Hansen, 2000). The temporal window for this FIR filter is determined individually (see Annex B on Hyper-parameters exploration), with distinct response functions trained for each pair of SoundNet features and brain regions. However, because the SoundNet features are non-causal, the temporal kernel may also account for differences in the temporal scales of the SoundNet features and fMRI brain activity, rather than solely modelling hemodynamic processes. In other words, the brain encoding layer simultaneously aligns artificial and biological neural processes over time while capturing hemodynamic processes.

#### Targets for brain alignment

The encoding layer was trained using two different brain targets, depending on what type of fMRI processed data the network learned to predict, thus yielding two different models:

- **STG model:** A model to predict fMRI signal from each 556 voxels located in the STG middle mask, at every TR, resulting in a prediction matrix’s size of 556 voxels prediction by the selected number of TR (see Annex Table 1 for the exact TRs number for every subject). We refer to this model as the *STG model* in the result section. This model is an evaluation of how well SoundNet predicts auditory fMRI activity in our settings, so we can estimate impact of brain alignment at the voxel level. For training data, we used data with or without spatial smoothing, to evaluate potential effects of spatial smoothing on encoding brain activity.
- **Whole-brain model:** A model to predict the average fMRI signal for all voxels of a parcel at every TR, resulting in a prediction matrix’s size of 210 parcels (also designed as ROI) prediction by the selected number of TR. We refer to this model as the *Whole-brain model* in the result section. The intention for this model is threefold (1) to verify which brain regions can be predicted by the model using audio as an input, (2) to check which ROIs are impacted by brain alignment and (3) to test whether individual variability has an impact on prediction performance and brain alignment.

#### Fixed training parameters

To train this architecture, we used AdamW (Loshchilov & Hutter, 2019) as an optimizer for L2 regularisation with weight decay, and we applied a learning rate scheduler that reduces the initial learning rate if no progress is achieved by the optimizer. The weight decay means that the brain encoding layer acts analogously to a Ridge regression, in effect regularising the regression parameters through shrinkage. MSE loss is used to minimise the difference between the predicted and actual fMRI signal.

For training individual models, we used the fMRI data from subjects watching the first three seasons of *Friends*. For each subject, we used 75 percent of the dataset for training, corresponding to 21 hours of training dataset. The remaining 25 percent was used for validation, corresponding approximately to 7h of the dataset, and we use all episodes of Season 4 (around 9h of audio) only for testing (see Table 2). In addition to individual models, we also trained group models to evaluate if a greater amount of training data could lead to better prediction results on one subject’s brain activity than using only fMRI data from the same subject. To fairly compare individual and group models, we decided to design a group model specific to each individual model: Unlike individual models where we used fMRI activity from one subject, we used fMRI activity from the other 5 subjects to train the corresponding group model (around 105h of dataset used for training, and 34h). With this method, group models performance won’t be influenced by individual features, and only by training dataset size.

**Table 2.**
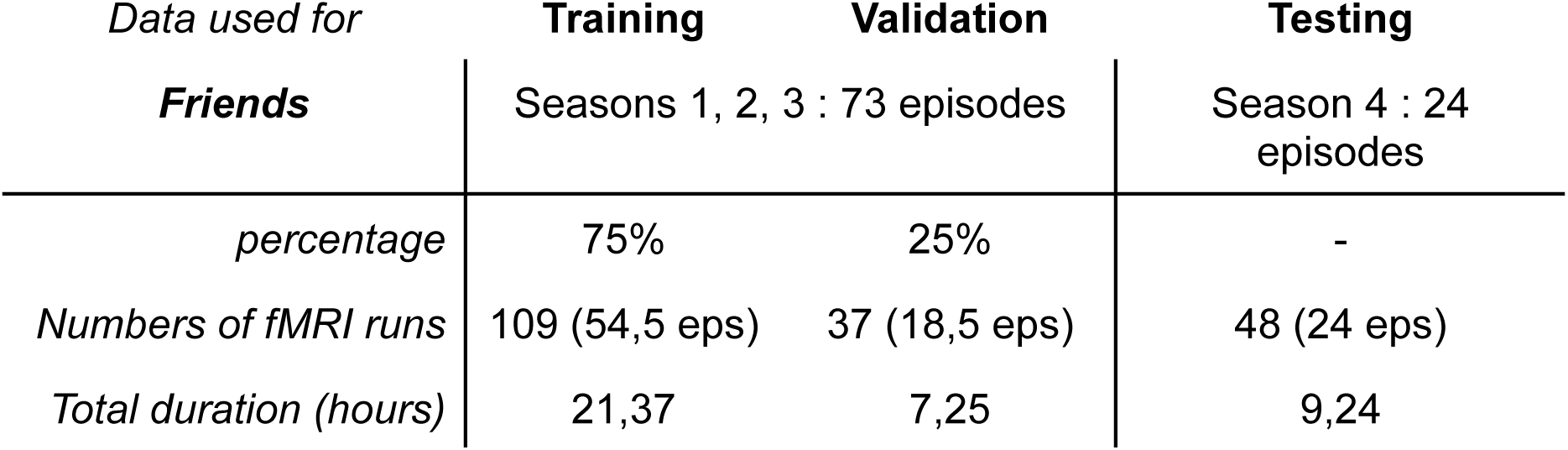
Repartition of the fMRI dataset *Friends* used for individual models between training, validation and testing. Each episode is split into two halves, with one half shown during an fMRI run.

| *Data used for*<br>***Friends*** | **Training** | **Validation** | **Testing** |
| --- | --- | --- | --- |
|  | Seasons 1, 2, 3 : 73 episodes |  | Season 4 : 24 episodes |
| *percentage* | 75% | 25% | - |
| *Numbers of fMRI runs* | 109 (54,5 eps) | 37 (18,5 eps) | 48 (24 eps) |
| *Total duration (hours)* | 21,37 | 7,25 | 9,24 |

To evaluate the accuracy of the model’s prediction, we computed the coefficient of determination r^2^ between the prediction and the corresponding fMRI time series for each region or voxel, for the entirety of the selected dataset (training or testing).

#### Hyper-parameters exploration

The goal of this study is to compare performance of an auditory AI network and of the same network but brain-aligned. We decided to realise the hyperparameter grid-search on the original trained SoundNet, with no fine-tuning, which we will consider as a fixed backbone, also referred to as the Baseline model. Through this grid-search, we were looking for an optimal set of hyperparameters to ensure SoundNet prediction performance as an encoding model as well as accounting for individual variability in the fMRI dataset. By going through these optimization steps, we have a better estimation of how much fine-tuning with brain representation impacts a network, for both brain encoding and network performance in classic AI tasks. The selected parameters to be tested only affect the training of the encoding layer at the end of the network.

For each individual model, we optimised different hyperparameters and criteria that could impact the final results (see Annex B and Annex D) :

- the duration of the audio waveforms given as input in each training iteration,
- the value of the learning rate at the beginning of the training,
- the size of the kernels in the encoding layer,
- the initial weight decay,
- the minimal change value considered for early stopping (referred to as “delta”),
- the number of epochs where such delta change is not present before stopping the training (“patience”).

For the corresponding group model, trained with data from the remaining five subjects, we used the median value of the results from all five individual models.

#### Fine-tuning the model

While only the encoding layer has been trained in the baseline model, the brain-aligned models have part of the original SoundNet’s parameters retrained to adjust prediction on individual fMRI data. As our architecture can be trained as an end-to-end network, we decided to test different levels of fine-tuning, from training only the last original layer of SoundNet (Conv7), to training the entirety of the network (Conv1). As such, we obtained 7 fine-tuned models both for Whole-brain and middle STG: Conv1, Conv2, Conv3, Conv4, Conv5, Conv6 and Conv7, each referring to the depth of the network that has been trained. Amongst these 7 models, we selected brain-aligned models that have the best ratio between brain encoding performance and training efficiency. We found that models where SoundNet has been fine-tuned up until Conv4 (referred as Conv4 models) achieve the best trade-off (see Annex E).

#### Models Comparison and statistical analysis for brain encoding

In order to evaluate the encoding performances of the baseline and brain-aligned models, we tried to predict fMRI activity with a null model, using the same architecture as the other models, but with randomly initialised weights. We used a Wilcoxon test (with a threshold of 0.05) to determine if the difference of the r^2^ value of a region / voxel between the null model and the baseline or brain-aligned model was significant across all 48 runs (half-episodes) of *Friends* season 4. As we repeated the same test for 210 regions or 556 voxels, we corrected the p-values obtained through the Wilcoxon test with a False Discovery Rate (FDR), using the Benjamini-Yekutieli procedure (Benjamini & Yekutieli, 2001), to take in account possible dependencies between tests derived at different regions / voxels. Only significant regions with a false-discovery rate *q* inferior to 0.05 were considered as significant. We repeated the same procedure to determine if the difference of r^2^ scores between baseline and fine-tuned models were significant, to evaluate if fine-tuning SoundNet on brain representations had an impact on SoundNet performances.

#### Identification and impact of audio annotations in the dataset

To understand the potential effect of brain alignment on the network’s performance, we analysed if changes in prediction could be driven by specific audio annotations present in the dataset used. Annotations were generated using a ResNet22 network pretrained on AudioSet (Gemmeke et al., 2017, Kong et al., 2020), a dataset including a large diversity of naturalistic audio sounds, ranging from *human voice* to *vehicle*, annotated through 527 labels with different categories and subcategories.

We segmented the audio track from every fMRI run (half episode) in 10 seconds audio excerpt, and used ResNet 22 to estimate the proportion of audio identified under each label for every excerpt. A subset of Audioset labels were aggregated into 8 annotations (see Annex Table 2 for details). For each run, we computed the average proportion of annotations across all 10 seconds excerpts, giving us in total 48 estimations of every annotation for each season of *Friends*.

To determine if the different seasons of *Friends* differ qualitatively and quantitatively, we computed a multivariable regression using the ordinary least square method (OLS) to assess if the proportion of audio labelled under each selected category was significantly different between each season (threshold for significativity set at 0.05). Finally, to evaluate if the difference in prediction accuracy between the baseline and brain-aligned models could be explained by the presence of specific annotations, we also did an OLS regression for each subject, using the difference in r^2^ score (maximum value amongst 210 ROI or 556 voxels) between the baseline and the brain-aligned individual models for each run of Season 4 as our dependant variable, and the difference from the mean proportion of each category for every run as our regressors (threshold for significativity set at 0.05). All statistical analyses have been done using the python library *Statsmodels*.

### Evaluating the models on HEAR

To evaluate how brain alignment impacted SoundNet performances, we tested every brain-aligned and baseline model on the Holistic Evaluation of Audio Representation (HEAR) benchmark (Turian et al., 2022). HEAR was proposed as an AI auditory competition during NeurIPS 2021, and gave the possibility to multiple research teams to test their network architectures and models. This benchmark has been made to evaluate how audio representations from a model are able to generalise over a series of 19 diverse audio tasks, including ESC50 and DCASE 2016, ranging from speech detection to music classification. A wide range of models have been evaluated with this API, resulting in a public ranking of auditory AI models in terms of transfer learning capability at the time of the competition (2022).

As some of the tasks required different inputs, the authors provided an API^5^ and preformatted datasets^6^ together with the evaluation code. We followed the API specifications, and extracted the representation of the Conv7 layer to use as scene embeddings for classification/labellisation tasks using the entire audio, and calculated timestamp embeddings (i.e. a sequence of embeddings associated with their occuring times) using the Conv5 layer, for sound event detection or transcription tasks. Both these embeddings are exposed to the HEAR API, which performs the evaluation of all 19 tasks, by using the embeddings as fixed input to train a downstream multi-layer perceptron (MLP). Depending on the task, the final layer could be softmax or a sigmoid, with cross-entropy loss. Details of the hyperparameters for the MLP training can be found in Appendix B of Turian et al. paper (2022).

For each task of the HEAR benchmark, we quantified the change in ranking for the SoundNet model before versus after fine-tuning with brain data, for each subject separately, and for each type of target (Full Brain vs STG). We applied a Wilcoxon test to determine an overall gain (or loss) in ranking across all the 19 tasks available in HEAR for each configuration separately.

## Results

### Comparing sound annotations in the training and test set

We first evaluated to what degree the sound distribution in our training set (*Friends* seasons 1 to 3) matched our test set (*Friends* season 4). For this purpose, we generated annotations of the sounds of each half-episode presented during a single fMRI run using a residual network ResNet 22, pretrained onAudioSet to label audio (Gemmeke et al., 2017, Kong et al. 2020). We further grouped these annotations into meta-categories and only selected the categories where at least 1 percent of the audio could be recognized within. These include categories such as *Music*, *Laugh*, *Women/Men speak* or *Applause*, with the category *Talking* being the most present amongst all (around 82% through all four seasons, see figure 3 for details). We then compared the distributions and found few statistically significant differences, and no substantial differences between seasons 1-3 and season 4. This result confirms that our generalisation experiment is a large-scale within distribution generalisation, at least in terms of these high level categories.

**Figure 3.**
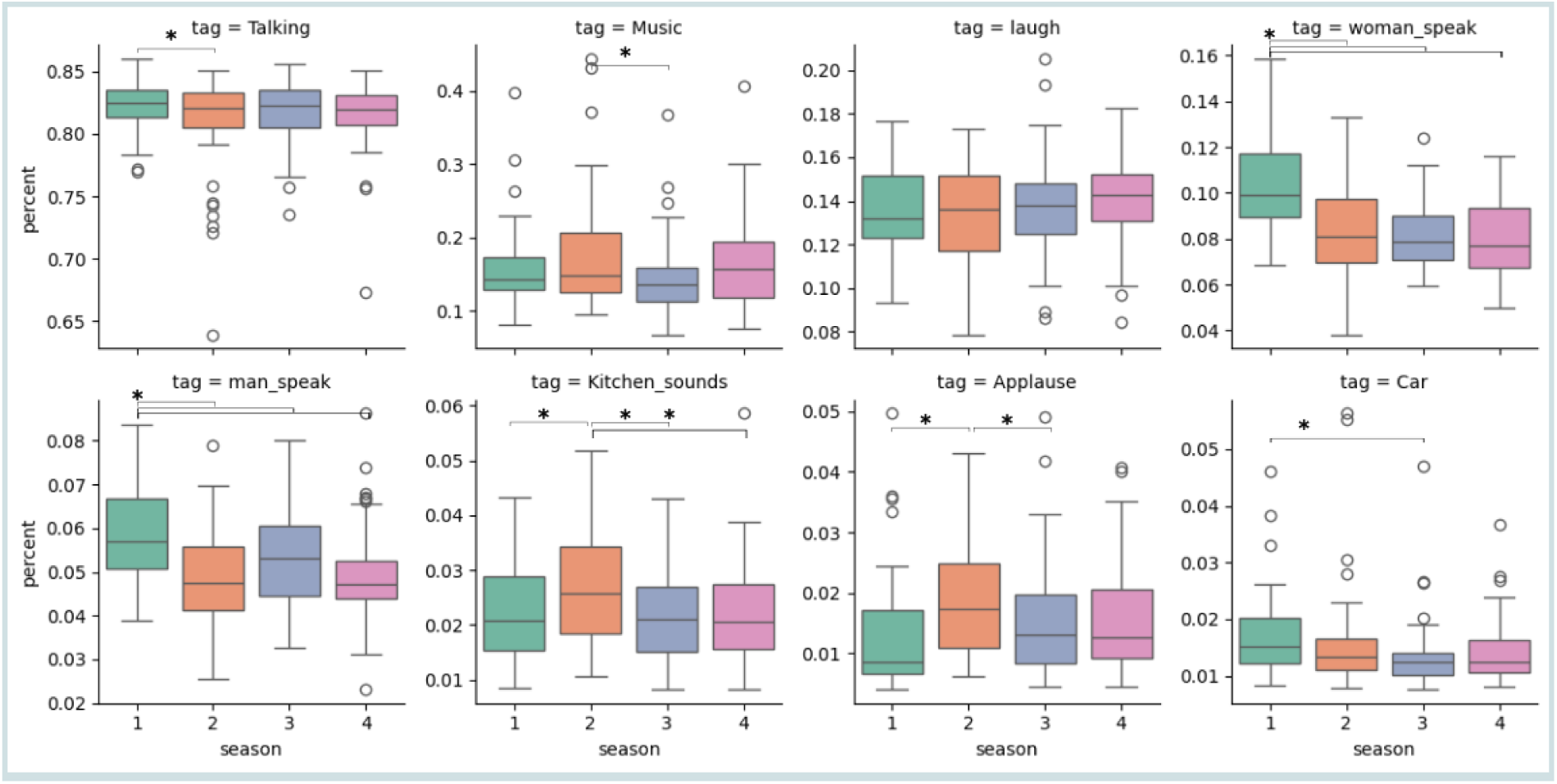
Proportion distribution of labelled audio in all half-episodes between *Friends*’s seasons. The proportion of labelled audio for each half-episode has been obtained using a ResNet 22 pretrained on AudioSet.

### Baseline brain encoding using pretrained SoundNet

#### SoundNet successfully encodes brain activity in the auditory cortex

We first tested the ability of our baseline model, SoundNet, to predict fMRI signals in different brain regions, using seasons 1-3 as training and validation and season 4 as test. It performed well, especially in the STG. Figure 4 shows almost all subjects had higher r² scores in the middle STG (STGm) than other regions (q < 0.05 for all subjects), except sub-05 whose best predicted parcel was the Middle Temporal Gyrus (MTG) superior. The posterior STG (STGp) also consistently ranked second in terms of prediction accuracy (see Annex Figure 2).

**Figure 4.**
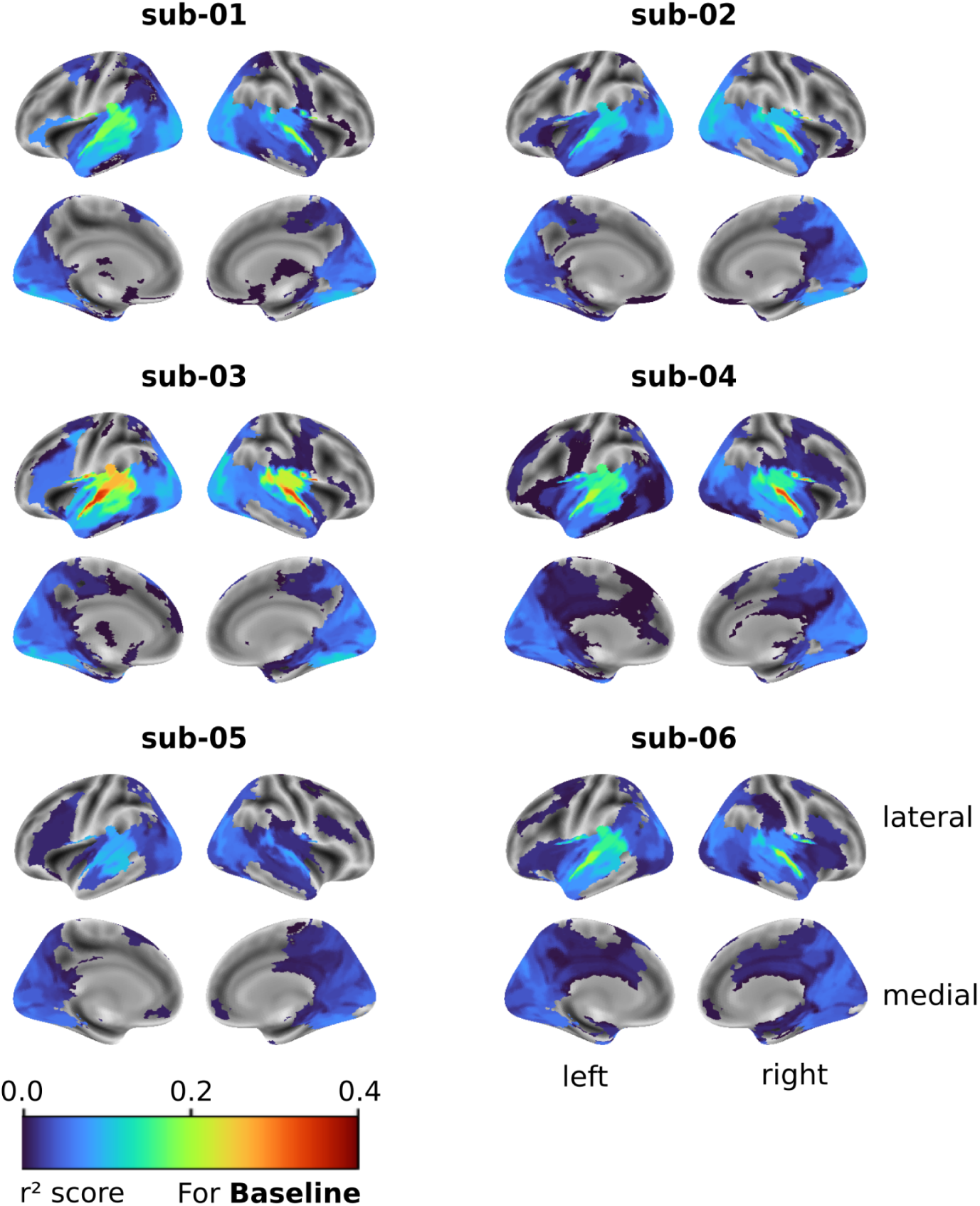
Full brain encoding using SoundNet with no fine-tuning. Surface maps of each subject, showing the r² value for all ROIs from the MIST ROI parcellation. Only parcels with r² values significantly higher than those of a null model initialised with random weights are shown (Wilcoxon test, FDR q < 0.05). Regions with highest r² scores are the STG bilaterally, yet significant brain encoding is achieved throughout most of the cortex, with relatively high values found in the visual cortex as well.

**Figure 5.**
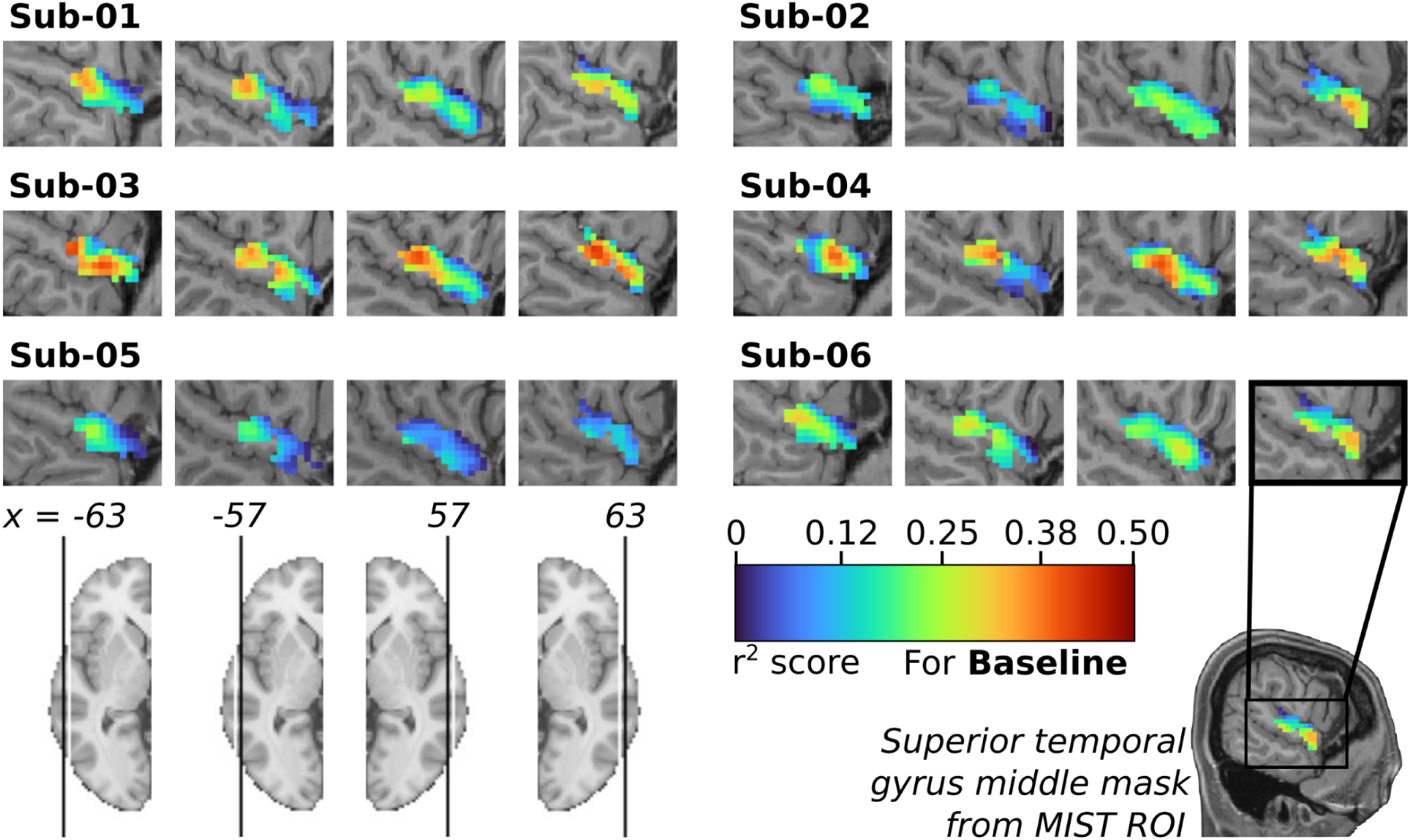
STG encoding using Soundnet with no fine-tuning and fMRI data with spatial smoothing. Mapping of the r² scores from 556 voxels inside the cerebral region defined as the Middle STG by the parcellation MIST ROI, computed by the individual Baseline model. To have a better representation of the STG, 4 slices have been selected in each subject, 2 from the left hemisphere (-63 and -57) and 2 from the right hemisphere (63 and 57). Only voxels with r² values significantly higher than those of a null model initialised with random weights are shown (Wilcoxon test, FDR q < 0.05). Individual anatomical T1 have been used as background.

SoundNet also accurately predicted other auditory regions like the MTG and Heschl’s gyrus in most subjects (Figure 4). This result supports our hypothesis that our baseline model can encode auditory processing from natural stimuli like movies, with some notable variations in performance between subjects, e.g. substantially higher performance was achieved in STG for sub-03 and sub-04.

SoundNet also encodes brain activity in the visual cortex and other regions Apart from the auditory cortex, brain activity in other regions was also predicted by the models; for most subjects we observed ROIs in the visual cortex such as the Lateral Visual Network DorsoPosterior (LVISnet_DP) and the Ventral Visual Network (VVISnet), respectively scoring as high as 0.12 and 0.11 (max scores in sub-03). These ROIs proved to be the best predicted ROIs after the STG and the MTG in Sub-01, Sub-02, Sub-05 and Sub-06, revealing that our baseline models were also able to encode aspects of the processing of an audio stimulus outside of the auditory cortex.

#### SoundNet encodes high resolution brain activity in the Superior Temporal Gyrus

When training the model to predict fMRI time series where a 5mm gaussian kernel has been applied for spatial smoothing, we found SoundNet could predict fMRI signals from voxels in the middle STG for all subjects. Most voxels’ fMRI activity was accurately predicted, with r² scores significantly different from the null model (514 to 555 significant voxels out of 556). Subjects 03 and 04 showed the best performance (average max r² of 0.45), while subject 05 performed worse (average max r² of 0.27). These results are consistent with the current literature regarding encoding activity from the auditory cortex and SoundNet ability to encode brain activity in the auditory cortex (Caucheteux et al., 2023, Nishida et al, 2020), and confirm that our model can predict fMRI activity linked to auditory processing at a high spatial resolution. It also shows similar individual differences as observed in full brain encoding. However, when using data with no spatial smoothing, we observe an important decrease in the number of voxels well predicted by the baseline model (see Annex Figure 3 and Discussion).

### Fine-tuning SoundNet with individual brain encoding

#### Individual models do not benefit of the brain alignment the same way

We next examined the fine-tuning impact on the brain-aligned models compared to the baseline models. After fine-tuning, the top-predicted parcels for Conv4 models were the same as those at baseline (Figure 6, left side of each subject panel). For most subjects, STGm and STGp remained the highest-scoring ROIs, with the exception of subject 02 model: while the right STGm is still best predicted, some visual cortices were better encoded than STG regions. We looked at which brain ROIs had the most improvement using Conv4 brain-aligned models.

**Figure 6.**
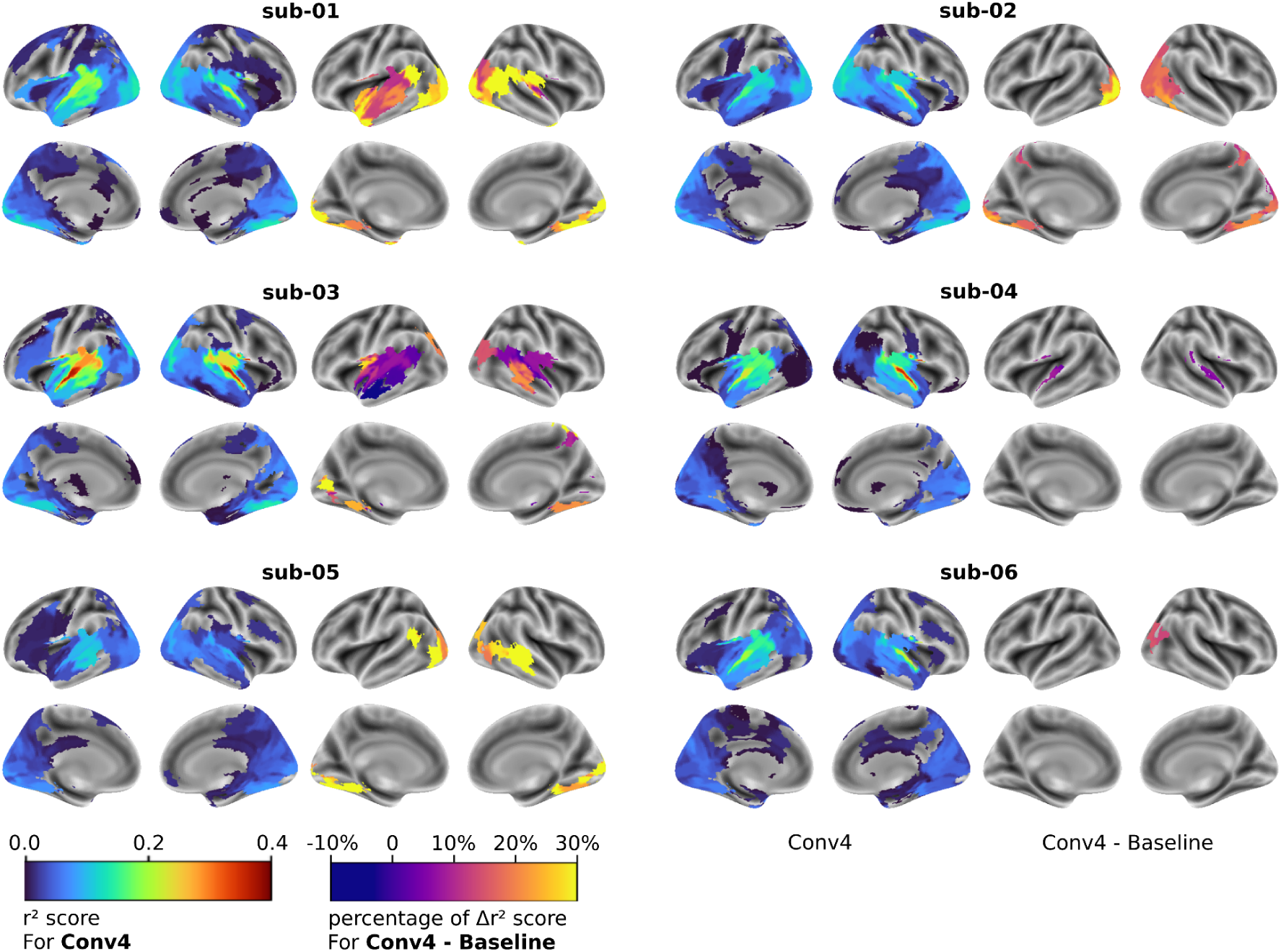
Individual impact of Brain-aligned SoundNet on the full brain encoding. For each subject, on the left side : Surface maps of the r² scores computed with each individual Conv4 model, for the 210 ROIs of the MIST ROI parcellation. Coloured ROIs have a r² score significantly greater than the null model (Wilcoxon test, FDR q <0.05). On the right side : surface maps of the percentage of difference in r² scores in each ROI between individual Conv4 and Baseline models. Only ROIs where Conv4 model have a r² score greater than +/- 0.05 and significantly greater or lesser than the baseline model are displayed (Wilcoxon test, FDR q <0.05)

We tested each ROI’s r² score for significant difference over the baseline and examined both the r² scores difference and the corresponding percentage of difference (see Figure 6, right side for a brain map of the percentage of r² score difference for each ROI, between the Conv4 and Baseline model for each subject): For most subjects, ROIs with the highest improvement in r² score gained between 0.01 and 0.04, with a relative gain from their original value of 15% to 30%, depending on the ROI and the subject. However, Sub-05 brain-aligned model, whose baseline model had the worse r² scores amongst all subjects, showed the highest relative gain in the MTG posterior (+ 0,03 r² score, corresponding to a gain of 167% of the original value). In general, ROIs with low r² scores (between 0.05 and 0.15) showed higher relative improvement than ROI with high r² score. The fine-tuning’s main improvements were not always in the auditory cortex: while for subjects 01, 04 and 05, the highest gain in r² score was in the right STG or MTG (between +0.02 and 0.04 r² score), for the remaining subjects it was located in the ventral, lateral or posterior visual network. Overall, fine-tuning improved the quality of brain encoding overall, with substantial variations across subjects both in terms of the magnitude and location of improvements.

#### Fine-tuning at the voxel level also leads to substantial improvements in brain encoding

We next wanted to see if fine-tuning also affected voxel-level fMRI signal predictions. We calculated the r² score difference between the baseline and the brain-aligned Conv4 model for each voxel in the STGm ROI, and only mapped those with significant differences (Figure 7). For both cases of training data (whether with or without spatial smoothing), we found voxels that were well predicted by the baseline models also had the most significant r² score increases for all subjects, although voxels with lower prediction accuracy were also impacted by fine-tuning. However, when using spatial smoothing, the median gain across all voxels was between 7 to 26% depending on the subject, lower than relative gains found in the brain-aligned models fine-tuned on the whole brain. While there were far less voxels being encoded when using data without spatial smoothing, the median gain in r2 score was higher for most subjects, between 10 to 115% (see Annex Figure 4). Overall, we found that fine-tuning improved brain encoding at the voxel level, with marked variations across subjects, and some departure from the impact of fine-tuning at the level of the full brain.

**Figure 7.**
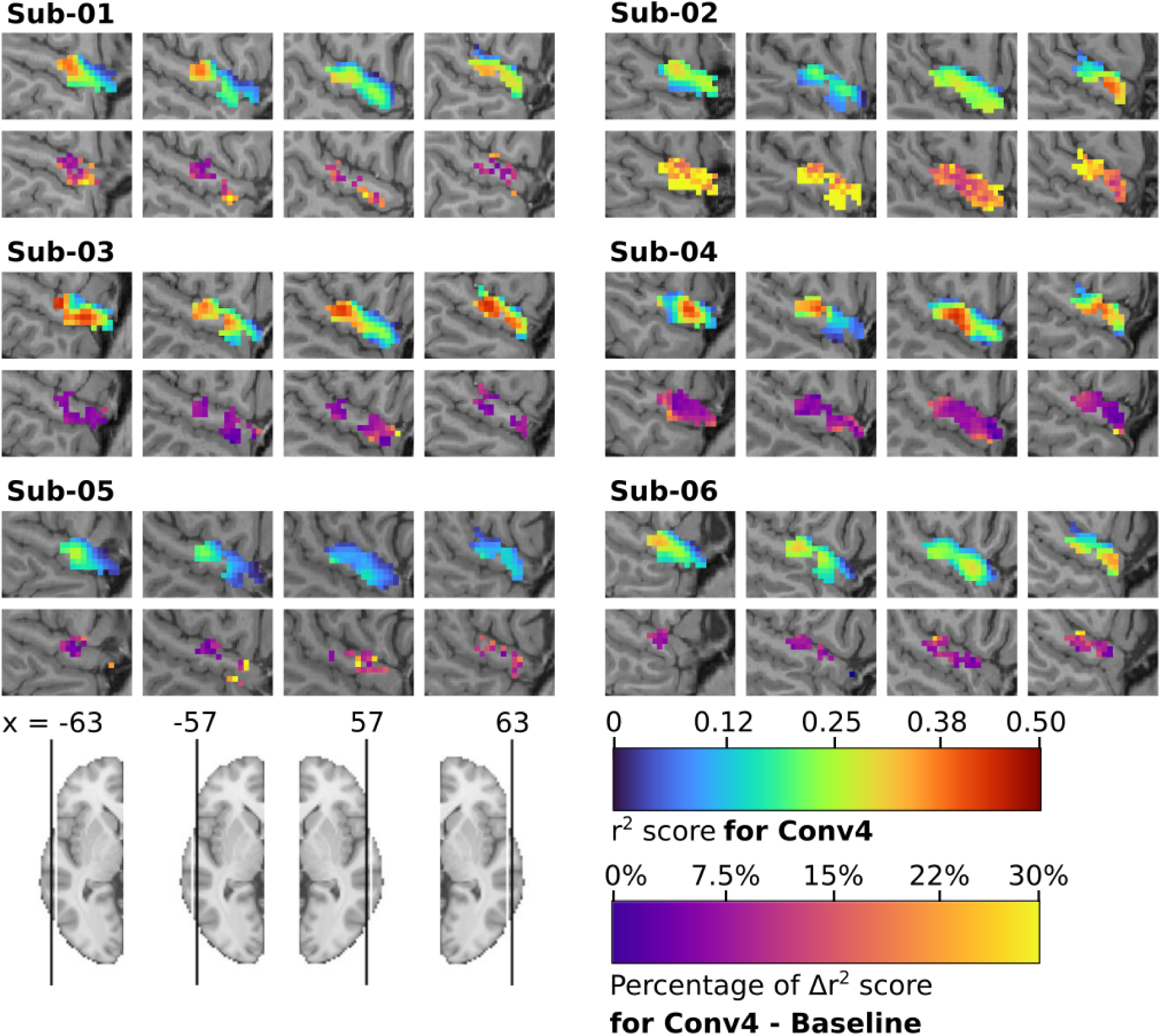
STG encoding using Brain-aligned SoundNet and fMRI data with spatial smoothing. For each subject, on the top : mapping of the r² scores from 556 voxels inside the cerebral region defined as the Middle STG by the parcellation MIST ROI, computed by the individual Conv4 model. Only voxels with r² values significantly higher than those of a null model initialised with random weights are shown (Wilcoxon test, FDR q < 0.05). For each subject, on the bottom : mapping of the difference of r² scores between the Conv4 model and the baseline model. Only voxels from the Conv4 model with r² values greater than +/- 0.05 and significantly greater or lesser than those of the baseline model are shown (Wilcoxon test, FDR q < 0.05). Individual anatomical T1 have been used as background.

#### Improvement in prediction in brain-aligned models does not associate strongly with specific audio features

To investigate some of the potential reasons that could lead to the prediction improvement observed with the brain-aligned model, we searched for correlation between the presence of specific features in the audio of season 4 and the prediction change. Using a residual network ResNet 22 pretrained on AudioSet to label the audio content, we computed the proportion of audio related to 28 categories for every half-episode of the season 4 of *Friends*. With the categories where at least 1 percent of the audio of season 4 could be associated with, we computed a multivariable regression using the ordinary least square method, to explain the difference in r² score for each half-episode in season 4. While tendencies can be found for categories such as *Talking*, *Kitchen sounds* or *Car*, no tag shows a significant impact through every model. This analysis did not reveal any major influence of the selected features on the prediction amelioration made by the brain-aligned models, but points to individual models being differently affected by the features, see Annex Figure 5 in Appendix H.

#### Brain-aligned individual models are subject specific, but group models show similar performance to individual models

Finally, we evaluated if the fine-tuned models were subject-specific, by applying models trained on data from one subject to fMRI signals collected on other subjects. When evaluating the difference for the maximal r² score in all roi (respectively all voxels) in the Whole Brain model (respectively STG model), we found that the model trained on one specific subject had the best performance to predict this specific subject’s data, both the Whole-Brain and middle STG models, with the exception of Sub-05 (figure 8). When looking at the difference in right STG middle ROI, sub-05 results are similar to other subjects. Overall, trained models appeared to exhibit subject-specific features.

**Figure 8.**
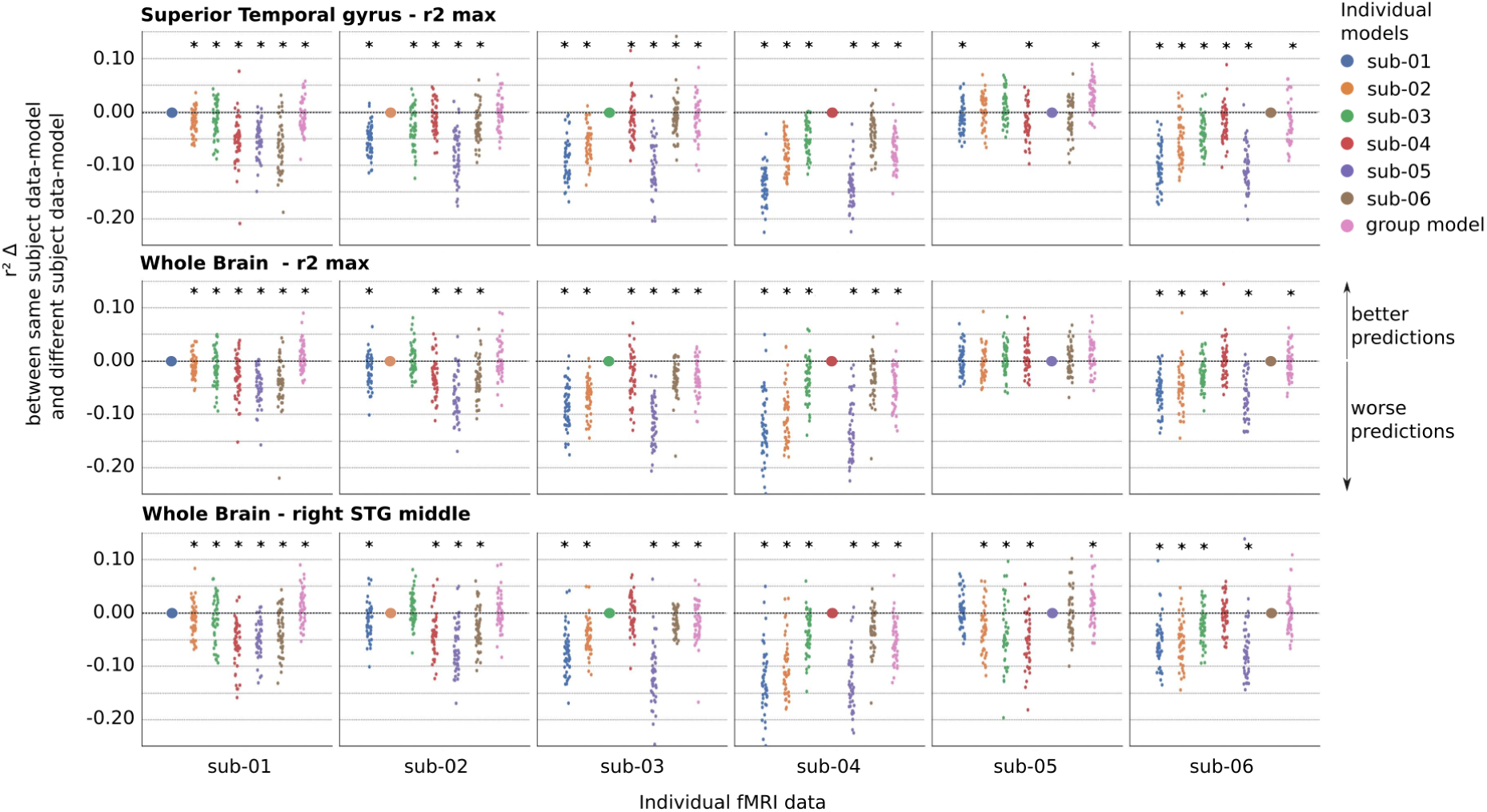
Comparison of prediction accuracy for subject-specific fMRI data using models trained on the same versus other subjects’ data. We computed the difference of r² scores computed by a brain-aligned model trained on data from the same subject as the test data, versus trained on one (from blue to brown) or a group of individual data (pink) different from the subject’s data used for testing. The difference is computed for each of the 48 half-episodes of the fourth season of *Friends*. A Wilcoxon test has been used to determine if the difference was significant between one individual model and the group model as well as each of the other 5 individual models (p < 0.05).

Results for group models show larger variability. For whole-brain models, two group models significantly outperform their respective individual models, two perform significantly worse, and the remaining two show no significant differences. For STG models, while 4 out of 6 individual models still significantly outperform their group counterparts, the group model performance often approaches that of the best encoding model.

Although the sample size is too small for definitive conclusions, it is noteworthy that subjects who benefited from a group model approach tended to have low baseline brain encoding performance.

### Fine-tuning improves SoundNet ranking in diverse AI auditory benchmarks

We aimed to evaluate the impact of brain alignment on the performance of SoundNet with downstream tasks, using the HEAR benchmark. For each task in the benchmark, we ranked both brain-aligned models and Soundnet amongst the 29 other models tested with this benchmark, and compared brain-aligned models against SoundNet. We analysed the difference of performance from three different angles:

- Between individuals and group models (12 models by task, including both Whole brain and middle STG, to evaluate if training on a larger dataset is more advantageous than training on individual dataset (see figure 9),
- Between Whole brain and middle STG models (6 models by task), to evaluate if the resolution of the training dataset could influence changes in performance in different tasks, (see figure 10),
- Between each individual subject (see figure 11), to see if individual features also have different effects depending on the task.

**Figure 9.**
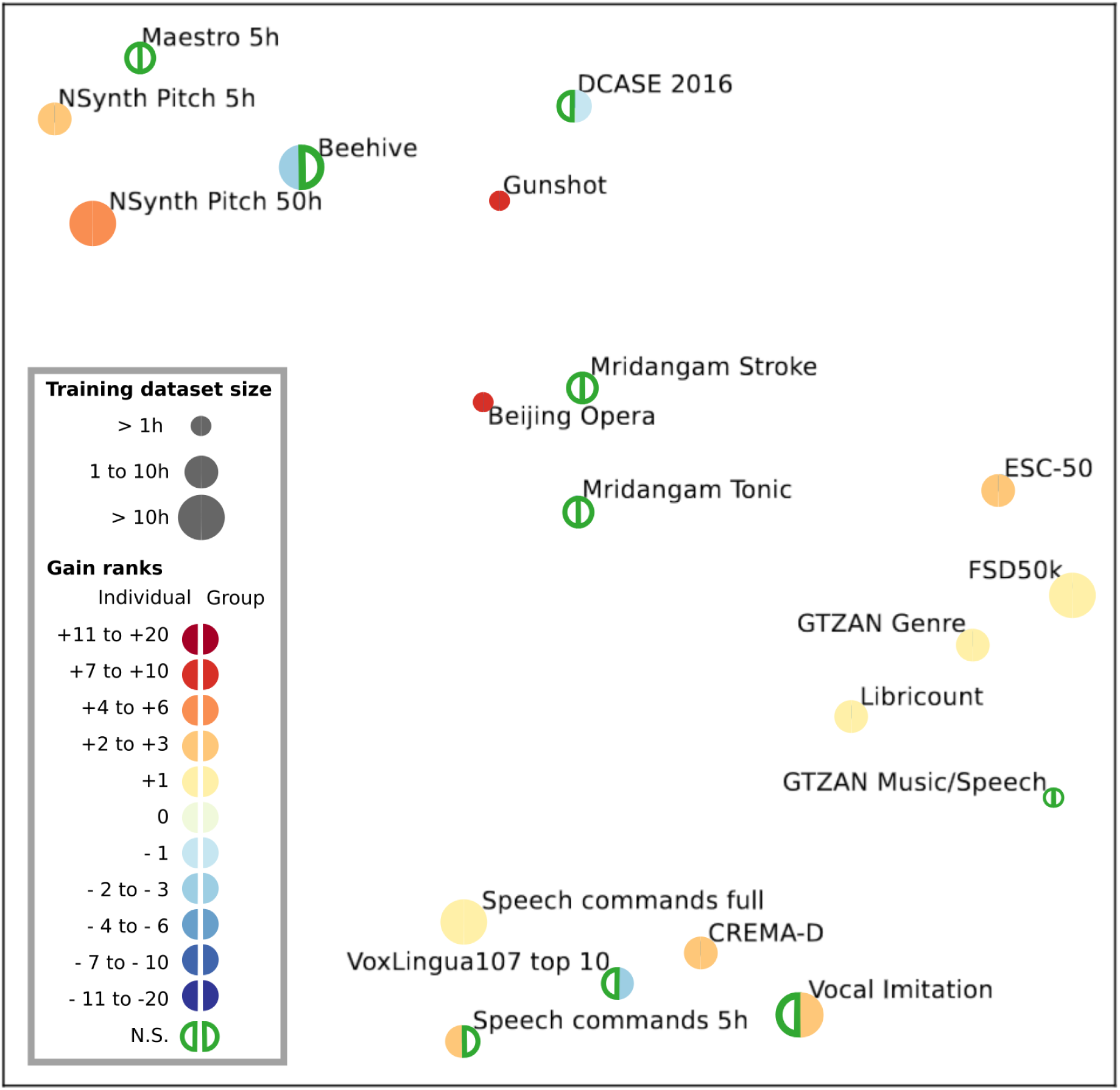
Rank variation between Conv4 and baseline models on all tasks from the HEAR benchmark. Adaptation of figure 2 of Appendix B from the original HEAR paper (Turian et al., 2022), showing a similarity visualisation of all 19 tasks, based upon normalised scores. For each task, the change of rank between the baseline model and the Conv4 model is symbolised by a coloured circle. Performance from both Whole-brain and STG versions of the individual models (half-circle on the left) and group models (half-circle on the right) have been averaged for each of the 19 tasks from the HEAR Benchmark. When the change of rank is equal to +1 (light yellow), Conv4 model is performing better than SoundNet at the task, but doesn’t outperform other models. Significativity has been tested using a Wilcoxon test (p < 0.05).

**Figure 10.**
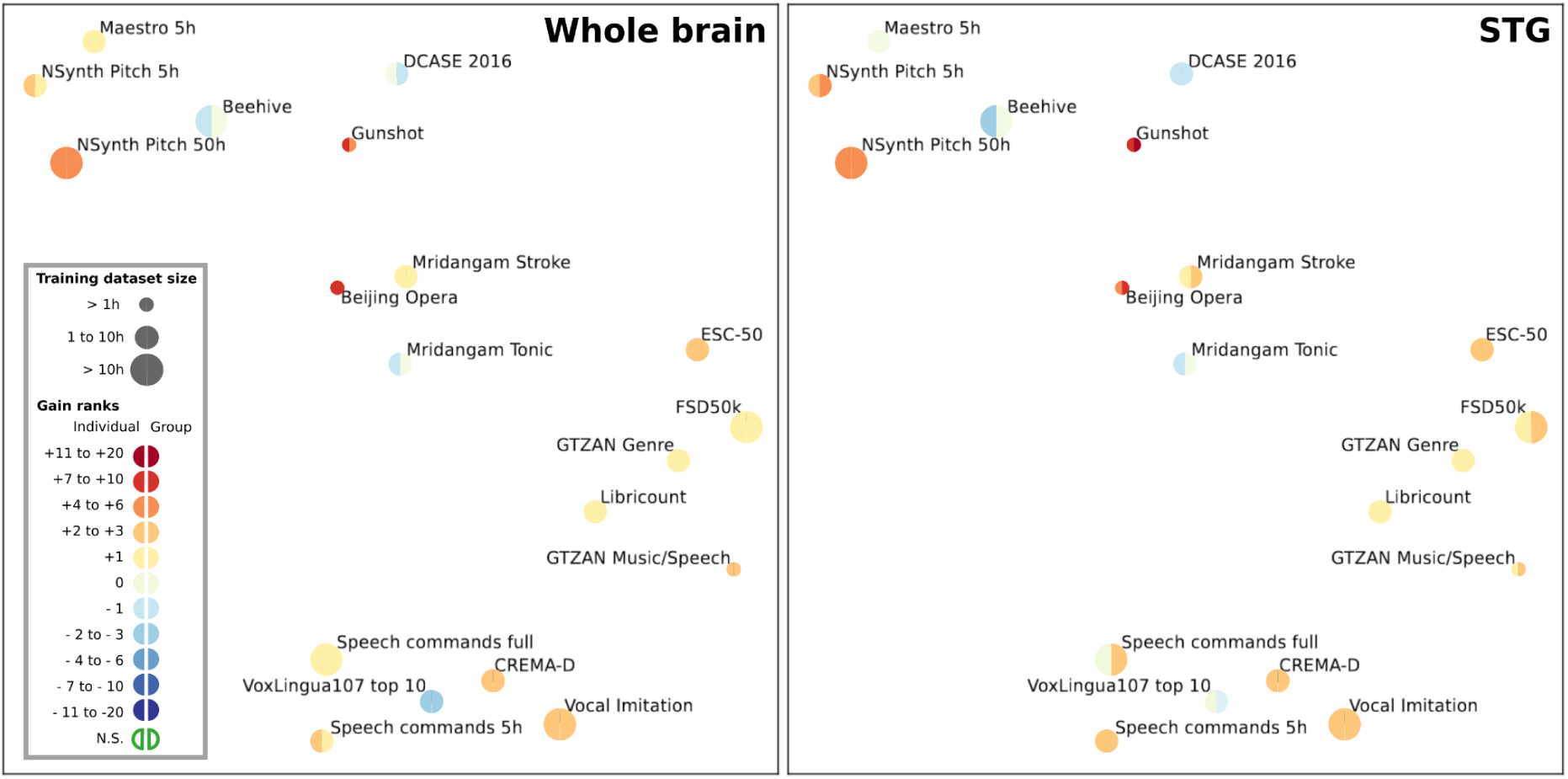
Rank variation between Whole brain and middle STG models on all tasks from the HEAR benchmark. Adaptation of figure 2 of Appendix B from the original HEAR paper (Turian et al., 2022), showing a similarity visualisation of all 19 tasks, based upon normalised scores. For each task, the change of rank between the baseline model and the Conv4 model is symbolised by a coloured circle. Left: Average change of rank with the Whole brain models (6 models for half a circle). Right: Average change of rank with the STG models (6 models for half a circle). Due to the low number of models per task, significance for each task hasn’t been tested at this level.

**Figure 11.**
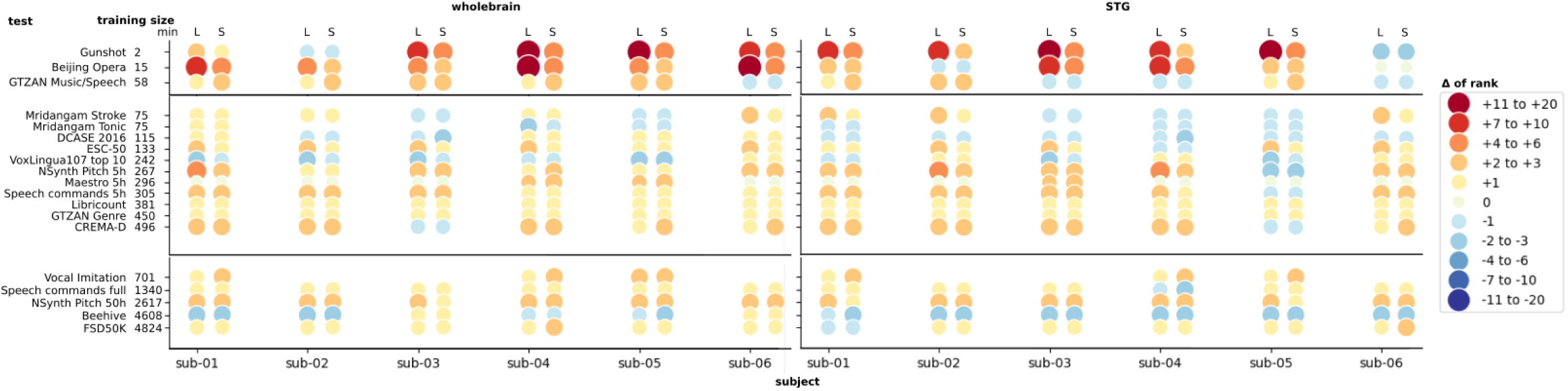
Rank variation between Conv4 and baseline models on all tasks from the HEAR benchmark, ordered by dataset size. Each individual Conv4 model (both Whole-brain and middle STG models) has been used to resolve the 19 tasks from the HEAR Benchmark, ordered by the size of training dataset available through the benchmark. We extracted from the official HEAR Benchmark Leaderboard^7^ the performances of 8 small models (up to 12M parameters) and 21 large models (from 22 to 1339 M parameters). We compared our brain-aligned and baseline models performances against the ones from large models (L columns, on the right side for each subject), and small models (S columns, on the left side for each subject). For each task, the change of rank between the baseline model and the Conv4 model is symbolised by a coloured circle.

#### Brain-aligned individual and group models display higher generalisation than SoundNet

After evaluating SoundNet in each task using the HEAR API, we found that SoundNet didn’t perform well in most tasks, being part of the least performing models multiple times. Brain-aligned models performed significantly better than SoundNet (p < 0.05) in 12 tasks out of 19, and performed worst in 2 tasks (DCASE 2016 and VoxLingua107 top 10) (see Figure 9 and Annex Figure 6 for more details). Gain in performance due to brain alignment is not related to a specific type of audio task, as brain-aligned models perform better in a variety of tasks, as shown in figure 10. When taking both middle STG and Whole brain models in account, we didn’t find any significant difference between individual and group models, both showing an average gain of 2 ranks.

We also observed that the brain alignment led to important gains in rank for a few tasks, such as Gunshot Triangulation, Beijing Opera and the NSynth Pitch (5h and 50h). Brain alignment seems to have the biggest impact with tasks involving small training datasets: For Gunshot Triangulation, brain-aligned models surpassed up to 18 others models, while they only had 2 minutes to retrain the parameters necessary to solve the task. Beijing opera shows similar results, but also has the highest standard deviation in ranking amongst all tasks. Considering that most models tested in this task scored around 0.95 in accuracy, we consider that the change in ranks observed for Beijing Opera is highly affected by the ranking distribution (see Annex Figure 6). In summary, brain alignment increases the generalisation capability of SoundNet.

#### Both whole-brain and voxel level fine-tuning results in better generalisation, but training on bigger datasets improve performance for voxel level models

Taking both individual and group models together, the comparison between whole-brain models and STG models yielded no significant differences in performance. Both showed an average improvement of 2 ranks across all tasks compared to SoundNet, with the highest rank gains observed in the same tasks: Gunshot and Beijing Opera.

However, when analysing separately the performance of group models and individual models, we found an influence of the resolution of the training dataset. At the voxel level, group models significantly outperformed individual STG models (p < 0.05), with an average rank gain for all tasks of 2.4 and 1.7, respectively. However, no significant differences were observed between individual and group models trained on the whole brain.

#### Performance of individual models vary between subject and task

For the individual models, we wanted to compare brain-aligned models performance with models of similar size (around 3 millions parameters). To do so, we divided the 29 models in two evaluation groups, depending on the number of parameters : a first group of 8 small models having less than 12 millions parameters, and a group with the remaining 21 models, ranging from 22 millions to 1339 millions parameters for the larger models (Figure 11).

Using Whole-brain fine-tuning, all 6 individual brain-aligned models displayed a significant gain in rank in the benchmark (over SoundNet and other models) (p < 0.05) (left panel of the figure 11). When comparing how brain-aligned models ranked amongst large models and small models (respectively *L* column and *S* column, for each individual model in figure 11), results were similar: all individual models had a significant increase in gain rank amongst small models, and 5 individual models displayed a similar increase amongst large models, with the exception of Whole-brain Sub-2 model, where its increase was still close to be significantly different (p = 0.057).

Results with individual middle STG models were more heterogeneous, with only sub-01, sub-02 and sub-03 Conv4 models ranking significantly higher than SoundNet, amongst small models and large models (right panel of Figure 11) Overall, all individual models show gains in rank in at least half of the tasks compared to SoundNet, but tasks with better performance are not the same through all individual models, showing important individual variability.

## Discussion

In this study, we explored the benefits of aligning a pretrained auditory neural network (SoundNet) with individual brain activity, both in terms of generalisation of brain encoding to new types of stimuli, and behavioural performance on a wide range of downstream tasks. Our results confirm substantial improvements in encoding brain activity, with gains extending beyond the auditory cortex, e.g. in the visual cortex. Importantly, brain alignment led to significant enhancements in performance across a broad range of auditory tasks when assessed using transfer learning. Our study also highlighted notable inter-individual variations, both in terms of the impact of brain alignment on brain encoding quality, and in the performance gains for downstream tasks.

### Can task-optimized ANN be aligned with brain activity ?

Our findings suggest task-optimised ANNs can successfully be aligned with individual brain activity. While we observed substantial enhancements in brain encoding quality, the extent of these improvements varied across both brain regions and individuals.

When models were directly fine-tuned to encode voxel-level STG activity with spatial smoothing, all participants exhibited modest but significant improvement in the superior temporal gyrus (STG) in most voxels. However, using data without spatial smoothing has reduced significantly the number of voxels encoded by both baseline and brain-aligned models, for all subjects. This result could be explained by a smaller tSNR at lower resolution, compared to data with spatial smoothing (Molloy et al., 2014). Considering the voxel size (2 mm isotropic), a 5 mm gaussian kernel can be considered as a good middle point between loss of spatial resolution and loss of tSNR. We also note that even without spatial smoothing, fine-tuning the models with fMRI data still improves the accuracy of the prediction in the few voxels that were well encoded in the baseline.

When models were fine-tuned on the entire brain, the STG remained the best predicted region in the brain after brain alignment. However, the impact of this process varied across both brain regions and individuals. For most subjects, regions outside the STG such as the visual cortex, experienced improvements comparable or greater to those in the STG. Different reasons can explain this result: Activation of the visual cortex by auditory stimuli have been observed in different contexts (Wu et al., 2007; Cate et al., 2009): Research in multimodal processing of spoken language found that visual cortices seem to be part of this specific process (Van Atteveldt et al., 2004; Seydell-Greenwald et al., 2023), and considering that the subjects were watching an audiovisual stimuli with a relative high amount of language content (around 80%), finding activation in the visual cortex in this context is coherent with the literature. To add up to this, the best predicted region in our study is the Superior Temporal Gyrus middle, always followed by the Superior Temporal Gyrus posterior and the Middle Temporal Gyrus, instead of the Heschl’s gyrus (Primary auditory cortex). These regions have also been shown to play a key role in the integration of visual and auditory information in the brain (Beauchamp et al., 2004; Van Atteveldt et al., 2004; Proverbio et al., 2011). Our results seem to indicate that our models have effectively learned the multimodal processing of audio stimuli within a naturalistic context, where the visual cortex is also involved. As the individual brain-aligned models are highly specific of the individual data used for the training, it is also possible that these results directly reflect specificity of individual processing learned with the brain alignment. However, it could also be due to the correlation of audio and visual features in our video stimuli (for instance, the presence of faces and lip movements during speech). It is challenging to draw direct comparisons with previous studies due to the sensitivity of the R2 metric to data acquisition and preprocessing decisions, including smoothing and voxel resolution.

### Impact of specific audio annotations on model’s performance

Further research is needed to understand the sources of variability in performance between individual models and to clarify which aspects of tasks and brain activity are critical for benefiting from brain alignment. Using a ResNet22 to annotate the audio content of Season 4 of *Friends*, we investigated potential audio features that might influence results. While no single feature consistently impacted all models, we observed tendencies indicating that individual models were influenced differently and weakly by various audio features. This observation is consistent with our downstream tasks (see below), which almost uniformly benefitted from brain alignment. Benefits of brain alignment thus do not appear to be limited to narrow categories of sound stimuli.

A limitation of this work is the temporal scale used for annotations: we averaged annotation results over half an episode, where multiple scenes with different audio contexts (e.g., kitchen, café, outdoors) occur together. Currently, we lack annotations related to the visual content of *Friends* episodes, which prevents us from encoding brain activity at the scene level. Investigating the correlation between brain encoding performance and audio annotations at the scene level, rather than over half an episode, would be an important step in further exploring these tendencies.

### Task performance on downstream tasks

We evaluated the performance of our brain-aligned models against SoundNet using the HEAR benchmark, which encompasses a variety of auditory tasks, and found that brain alignment generally benefited performance on downstream tasks. Few studies have employed a downstream task benchmark after brain alignment. Palazzo et al. (2020) reported modest performance gains in vision tasks post-alignment with EEG data. Nishida et al. (2020) reported similar findings with their audiovisual tasks, using fMRI, as well as Moussa et al. (2024) for semantic tasks. However, Schwartz et al. (2019) reported no significant change in performance after brain alignment with fMRI or MEG data. Our research differs notably in stimulus nature (a TV show), and the very large volume of fMRI data used for fine-tuning. While our results seem to align with the first three studies in terms of finding mostly moderate improvements in performance, this study is the first to examine a wider range of auditory downstream tasks. The primary goal of the HEAR benchmark is to evaluate the capacity of a network’s internal representations to generalise to new tasks with data of a different nature than what has been used to initially train the network. Considering this goal, brain-aligning a pretrained CNN network led to more generalisable representations, but also identify possibly large gains for downstream tasks with limited training data available: The two tasks that benefited the most are gunshot triangulation (a classification task) and Beijing Opera percussion (an instrument recognition task), which are small scale datasets (Turian et al., 2022) (training data corresponds to approx. 100s and 900s, respectively). However, our results also show improvements on much larger datasets, such as NSynth pitch classification on 5h and 50 hours of data (Engel et al., 2017), as well as modest benefits on a very large and difficult benchmark, FSD50k, a multi-label audio tagging dataset with more than 80 hours of training data (Fonseca et al., 2021). Taken together, the ability of our brain-aligned representations to generalise to small and larger scale datasets suggest both that they are general enough to generalise with few data, and flexible enough to enable gains on larger scale tasks.

### Do models benefit more from individual datasets, compared to bigger datasets ?

We compared performance of ANNs trained on individual datasets versus trained on bigger group datasets for both brain encoding (within distribution) and tasks from the HEAR benchmark (out of distribution).

For models trained to predict STG fMRI activity at the voxel level, we observed that 4 out of 6 individual models significantly outperformed group models trained on multiple subjects in predicting fMRI activity for the same subject. Among the remaining two, the group model for Sub-05 performed better than its corresponding individual STG model. Notably, the Sub-05 individual model was the weakest performer among all individual models, and the Sub-05 dataset had the lowest temporal signal-to-noise ratio (tSNR) of all datasets, see Annex B for more details. This suggests that, at the voxel level, having data with a high tSNR may be crucial for an individual model to capture subject-specific processing and outperform group models in predicting fMRI activity. However, it remains unclear whether individual specificity aids generalization to new tasks, as individual STG models performed worse than their respective group STG models on these tasks.

For whole-brain models, the benefits of individual models compared to group models remain unclear. While two individual models significantly outperformed their group counterparts in encoding brain activity, two performed worse, and no significant differences were observed for the remaining two. Additionally, group models do not show improved generalization, as individual and group models perform similarly on many downstream tasks. Given the current focus in the AI field on reducing computational costs and the challenges of acquiring sufficiently large fMRI datasets for training neural networks, this work suggests that smaller, individual fMRI datasets can be as effective as larger datasets in improving network generalizability for whole-brain models.

### Longer temporal windows for training led to better encoding results

At a technical level, hyper-parameter optimization was an important step for effective fine-tuning of SoundNet. We adopted an approach that combines multiple steps, starting with an extensive grid search on one subject, and then refining the optimal parameters per subject on a subset of the grid and parameters that had the most notable impact in our initial investigation (see Supplementary Results 1). A surprising finding was that the optimal duration of the input window for sound waves extended up to 70 TRs (105 sec). Two main factors may explain this observation. First, it is known that auto-regressive models of fMRI activity improve their performance even for very context windows. Our group recently published a study using the *Friends* dataset where we found the best model to use 286 seconds (4min and 46s) of fMRI data to predict the next time point (Paugam et al., 2024). There is thus evidence of long term memory processes in fMRI brain data. It is possible that these processes reflect in part exogenous stimuli such as sound. Another possibility is that SoundNet was trained to generate visual annotations from the sound of short videos. It is thus intrinsically biassed towards the duration of these videos, ranging from a few seconds to a few minutes. In any case, a take-away of our study is that brain alignment is sensitive to hyper-parameter optimization and this work may provide a guide for selecting ranges and parameters in future works.

### Problems of a Within-Distribution testing dataset for brain encoding

The annotation study showed that the TV show used in the dataset for this study presents strong similarities between each season, resulting in within distribution training and testing. This kind of training may possibly lack in diversity to pretrain an ANN and check for broad generalisation of representations. The CNeuroMod data collection includes a variety of stimuli beyond the *Friends* TV show, and it would be possible to check how different types of stimuli impact generalisation to downstream AI tasks. Additionally, we’re interested in investigating whether brain-aligned models lead to human-like similarity judgement patterns (Bakker et al., 2022).

### Limitations

A limitation of this study is to focus on a single pretrained network, Soundnet. Considering recent advances in AI auditory models performance (Schmid, Koutini, & Widmer, 2023), it would be important to study other architectures as well in the future, to evaluate if and how brain alignment impact could differ depending on the architecture used (e.g. Transformers versus convolutional networks), the number of model parameters and the type of data used for pretraining. We also found that SoundNet had a lower score in benchmarks such as ESC-50 and DCase, compared to the scores of its original paper. While we tried to stay as close as the original implementation, multiple reasons could explain this difference of score: it is possible that the original SoundNet paper used different embeddings to evaluate the model, compared to the embedding required by the HEAR benchmark. We also used Python with Pytorch to implement the brain-aligned models and end-to-end training, while originally SoundNet was done in Lua with Tensorflow. The conversion from one library to another could also have an impact.

It should finally be noted that the parcels used from the MIST ROI parcellation were based on non-linear alignment; while the models trained on the Whole-brain fMRI activity best predicted the STG, they also displayed important individual differences. We can’t exclude the possibility that specific ROIs in the auditory cortex and the visual cortex could be slightly misaligned with individual anatomy, which could partially impact the results.

### Conclusions

In our study, we developed the first set of auditory deep artificial networks fine-tuned to align with individual participants’ brain activity. This was made possible by the Courtois NeuroMod project’s massive individual data collection effort. We successfully fine-tuned a pretrained network called SoundNet to better encode individual participants’ brain signals, showing varying degrees of improvement over a model that only adds an encoding layer to predict brain signals. These brain-aligned models also improved in performance a pretrained network, trained without brain data on a diverse set of AI audio tasks, ranging from classifying pitch to determining the number of speakers. The brain-aligned models also demonstrate high potential for tasks with limited dataset available and few-shot learning. These findings open many avenues for future research, ranging from studying inter-individual variations to testing brain alignment for various model architectures, types of training data and types of downstream tasks.

## Supporting information

Appendices - Methods & Results

## Ethics

All subjects provided informed consent to participate in this study, which was approved by the ethics review board of the “CIUSS du centre-sud-de-l’île-de-Montréal” (under number CER VN 18-19-22).

## Data and Code Availability

The fMRI data used to train the model is openly available through registered access at link https://www.cneuromod.ca/access/access/. All the code used to train the models, produce the results and figures are freely accessible on a github repository at https://github.com/brain-bzh/cNeuromod_encoding_2020.

## Authors Contributions

Prototyping, development, and fine-tuning of the models have been done by MF and NF, with advice from PB.

API development to adapt models for evaluation with the HEAR benchmark has been done by NF, LT and MLC.

Statistical analysis and visualisation has been done by MF, with advice from NF and PB.

Manuscript has been written by MF and PB, with additional reviewing and contributions on the methods and discussion section by NF. The AI tool ChatGPT4 has been used only to help in improving text architecture and writing style.

## Funding and Acknowledgments

The Courtois project on neural modelling was made possible by a generous donation from the Courtois foundation, administered by the Fondation Institut Gériatrie Montréal at CIUSSS du Centre-Sud-de-l’île-de-Montréal and University of Montreal. The Courtois NeuroMod team is based at “Centre de Recherche de l’Institut Universitaire de Gériatrie de Montréal”, with several other institutions involved. See the cNeuroMod documentation for an up-to-date list of contributors (https://docs.cneuromod.ca).

The work was partly supported by a grant from the Brittany region in France “Allocation de Recherche Doctorale” to NF, and a Digital Alliance Canada resource allocation grant to PB to access the Béluga high performance computing infrastructure. PB is supported by a salary award as senior fellow (chercheur boursier senior) of the “Fonds de Recherche du Québec - Santé”.

## Appendices

### Appendix Methods

#### Annex A fMRIprep workflow

Results included in this manuscript come from preprocessing performed using *fMRIPrep* 20.2.5 (@fmriprep1; @fmriprep2; RRID:SCR_016216), which is based on *Nipype* 1.6.1 (@nipype1; @nipype2; RRID:SCR_002502).

A total of 2 T1-weighted (T1w) images were found within the input BIDS dataset for each subject. Anatomical preprocessing was reused from previously existing derivative objects.

For each of the BOLD runs found per subject (across all tasks and sessions), the following preprocessing was performed. First, a reference volume and its skull-stripped version were generated by aligning and averaging 1 single-band reference (SBRefs). A B0-nonuniformity map (or fieldmap) was estimated based on two (or more) echo-planar imaging (EPI) references with opposing phase-encoding directions, with 3dQwarp @afni (AFNI 20160207). Based on the estimated susceptibility distortion, a corrected EPI (echo-planar imaging) reference was calculated for a more accurate co-registration with the anatomical reference. The BOLD reference was then co-registered to the T1w reference using bbregister (FreeSurfer) which implements boundary-based registration [@bbr]. Co-registration was configured with six degrees of freedom. Head-motion parameters with respect to the BOLD reference (transformation matrices, and six corresponding rotation and translation parameters) are estimated before any spatiotemporal filtering using mcflirt [FSL 5.0.9, @mcflirt]. First, a reference volume and its skull-stripped version were generated using a custom methodology of fMRIPrep. The BOLD time-series were resampled onto the following surfaces (FreeSurfer reconstruction nomenclature): fsaverage. The BOLD time-series (including slice-timing correction when applied) were resampled onto their original, native space by applying a single, composite transform to correct for head-motion and susceptibility distortions. These resampled BOLD time-series will be referred to as preprocessed BOLD in original space, or just preprocessed BOLD. The BOLD time-series were resampled into standard space, generating a preprocessed BOLD run in MNI152NLin2009cAsym space. First, a reference volume and its skull-stripped version were generated using a custom methodology of fMRIPrep. Grayordinates files [@hcppipelines] containing 91k samples were also generated using the highest-resolution fsaverage as intermediate standardised surface space. Several confounding time-series were calculated based on the preprocessed BOLD: framewise displacement (FD), DVARS and three region-wise global signals. FD was computed using two formulations following Power (absolute sum of relative motions, @power_fd_dvars) and Jenkinson (relative root mean square displacement between affines, @mcflirt). FD and DVARS are calculated for each functional run, both using their implementations in Nipype [following the definitions by @power_fd_dvars]. The three global signals are extracted within the CSF, the WM, and the Whole-brain masks. Additionally, a set of physiological regressors were extracted to allow for component-based noise correction [CompCor, @compcor]. Principal components are estimated after high-pass filtering the preprocessed BOLD time-series (using a discrete cosine filter with 128s cut-off) for the two CompCor variants: temporal (tCompCor) and anatomical (aCompCor). tCompCor components are then calculated from the top 2% variable voxels within the brain mask. For aCompCor, three probabilistic masks (CSF, WM and combined CSF+WM) are generated in anatomical space. The implementation differs from that of Behzadi et al. (2007) in that instead of eroding the masks by 2 pixels on BOLD space, the aCompCor masks are subtracted a mask of pixels that likely contain a volume fraction of GM. This mask is obtained by dilating a GM mask extracted from the FreeSurfer’s aseg segmentation, and it ensures components are not extracted from voxels containing a minimal fraction of GM. Finally, these masks are resampled into BOLD space and binarized by thresholding at 0.99 (as in the original implementation). Components are also calculated separately within the WM and CSF masks. For each CompCor decomposition, the k components with the largest singular values are retained, such that the retained components’ time series are sufficient to explain 50 percent of variance across the nuisance mask (CSF, WM, combined, or temporal). The remaining components are dropped from consideration. The head-motion estimates calculated in the correction step were also placed within the corresponding confounds file. The confound time series derived from head motion estimates and global signals were expanded with the inclusion of temporal derivatives and quadratic terms for each [@confounds_satterthwaite_2013]. Frames that exceeded a threshold of 0.5 mm FD or 1.5 standardised DVARS were annotated as motion outliers. All resamplings can be performed with a single interpolation step by composing all the pertinent transformations (i.e. head-motion transform matrices, susceptibility distortion correction when available, and co-registrations to anatomical and output spaces). Gridded (volumetric) resamplings were performed using antsApplyTransforms (ANTs), configured with Lanczos interpolation to minimize the smoothing effects of other kernels [@lanczos]. Non-gridded (surface) resamplings were performed using mri_vol2surf (FreeSurfer).

Many internal operations of *fMRIPrep* use *Nilearn* 0.6.2 [@nilearn, RRID:SCR_001362], mostly within the functional processing workflow. For more details of the pipeline, see the section corresponding to workflows in *fMRIPrep*’s documentation^8^.

The above boilerplate text was automatically generated by fMRIPrep with the express intention that users should copy and paste this text into their manuscripts *unchanged*. It is released under the CC0 license

#### Annex B Exploration of Hyperparameters space

**Annex Table 1.**
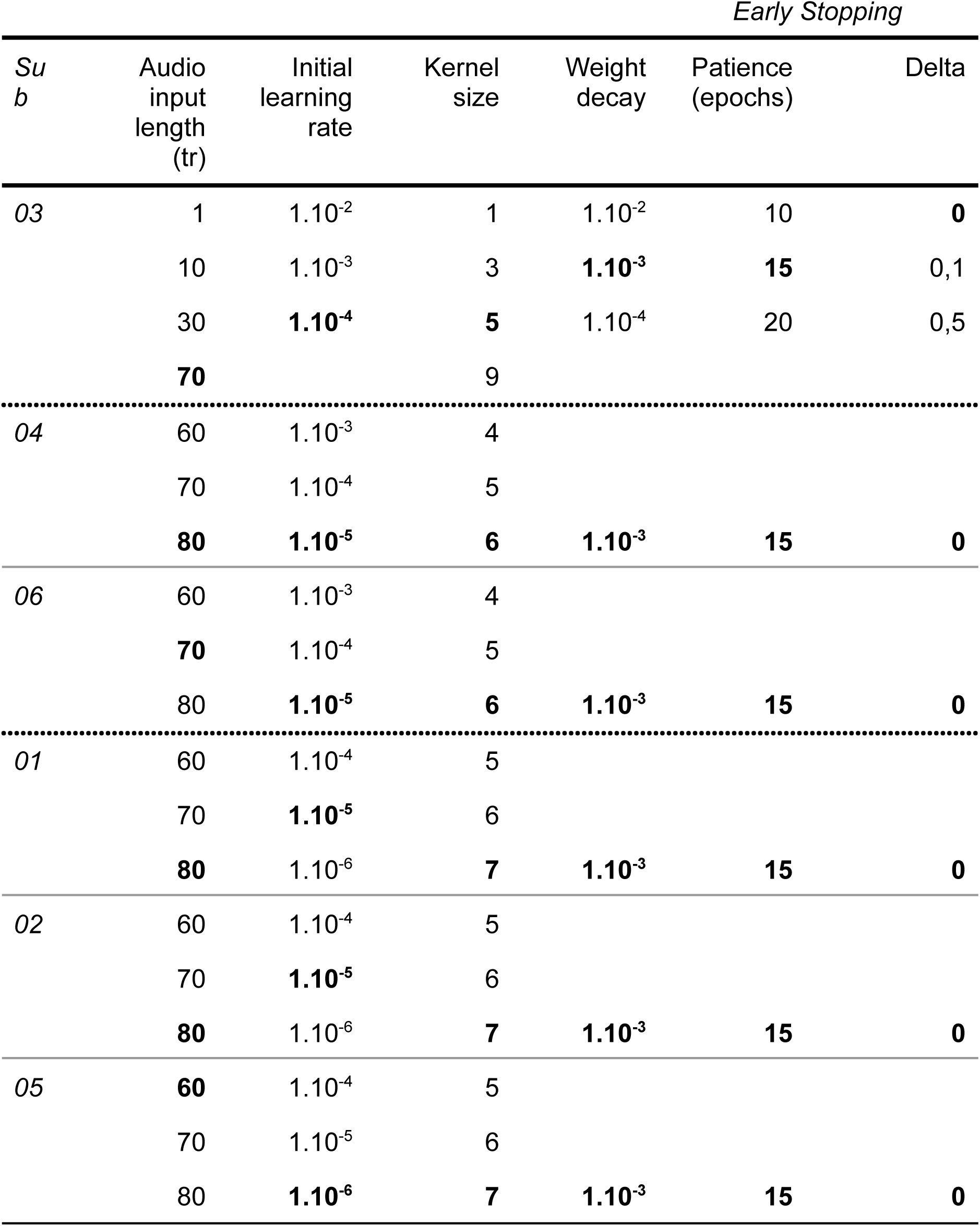
Hyperparameters values explored for subjects’ baseline model. The values selected for fine-tuning the models are marked in bold. The subject order has been constrained by the availability of all four season fMRI data for each subject at the time of the analysis, as this study was taking place in parallel to the data collection. For the kernel size and the learning rate, the range of values shifted by 1 unit between sub-06 and sub-01, as the optimal values for sub-4 and sub-06 appeared to be the maximal value of the explored window, and we wanted to ensure optimal hyperparameters for each subject.

As we are working on individual datasets, we wanted to explore the hyperparameters space for each subject’s baseline models. However, doing so would be particularly time-consuming and highly costly in terms of computational resources. In order to constrain the hyperparameters grid search, we decided to explore all parameters only with the baseline model trained on one subject fMRI dataset. We chose to use Sub-03, as their time series had the highest temporal Signal-to-Noise Ratio (tSNR) and lowest motion levels amongst all 6 subjects (Boyle et al., 2023). The tSNR has been averaged in the brain mask of each run of Friends seasons 1 and 2, as well as two others datasets of the Courtois Neuromod project. The tSNR has been extracted by MRIQC, using the MRIQC API (Esteban et al., 2017).

We trained the baseline model (fixed-weights SoundNet + encoding layer) to predict fMRI activity from sub-03, with 1296 different configurations of hyperparameters (see Table 3 for the selected values explored for each criteria). This step has been done twice, for sub-03 STG model and Whole-brain model, but as results were equivalent between both, we decided to only keep results from the Whole-brain model. To distinguish between all configurations of hyperparameters, we ranked trained models by their configuration and their prediction performance on the validation set: we used the r^2^ score of the best predicted parcel as our measure or performance. We computed a correlation matrix between ROI predictions from the best 100 configurations, to determine if some of the models trained on these configurations shared a similar parcel prediction pattern. We observed 2 to 3 clusters of configurations, so in order to better define these clusters, we used an agglomerative hierarchical clustering, with a linkage function computing the Euclidean distance between centroid of clusters, and divided the output in 3 clusters (UPGMC algorithm, as implemented by the Scipy library^9^). For each cluster, we computed the corresponding predicted brain map by averaging the prediction output of all configurations in the cluster. We then compared all 3 clusters, and chose the cluster with the highest maximal r^2^ score (best predicted parcel amongst 210) and highest mean r^2^ score (mean of all 210 r^2^ scores). When looking at the values for each hyperparameter, we determined which value was prevalent in the best cluster, by computing the statistical mode on the values selected for all configurations present in this cluster (see Supplementary Result 1).

After deciding which parameters and values had the most impact on the capacities of the baseline model to predict fMRI signals from sub-03 dataset, we switched to other subjects, this time exploring only the most impactful hyperparameters on sub-03 and a limited set of values around the optimal value found for sub-03 (see Table 3). As a result, we explored only 27 configurations for each of the remaining subjects. We defined the best hyperparameters values by computing the mode amongst the 10 most performing configurations. We did not use cluster analysis for other subjects, as we only had 27 configurations. As the data collection was done in parallel to the hyperparameter grid-search, we started this process with data from a few subjects (sub-03, sub-04 and sub-06). After selecting the hyperparameters configuration for both sub-06 and sub-04, we decided to adjust the window of tested values of two hyperparameters for the remaining subjects, as their optimal values seemed to always be the highest value for both sub-06 and sub-04 (see Supplemental Table 3 for more details).

#### Annex C Details of AudioSet categories used for audio annotation

**Annex Table 2.**
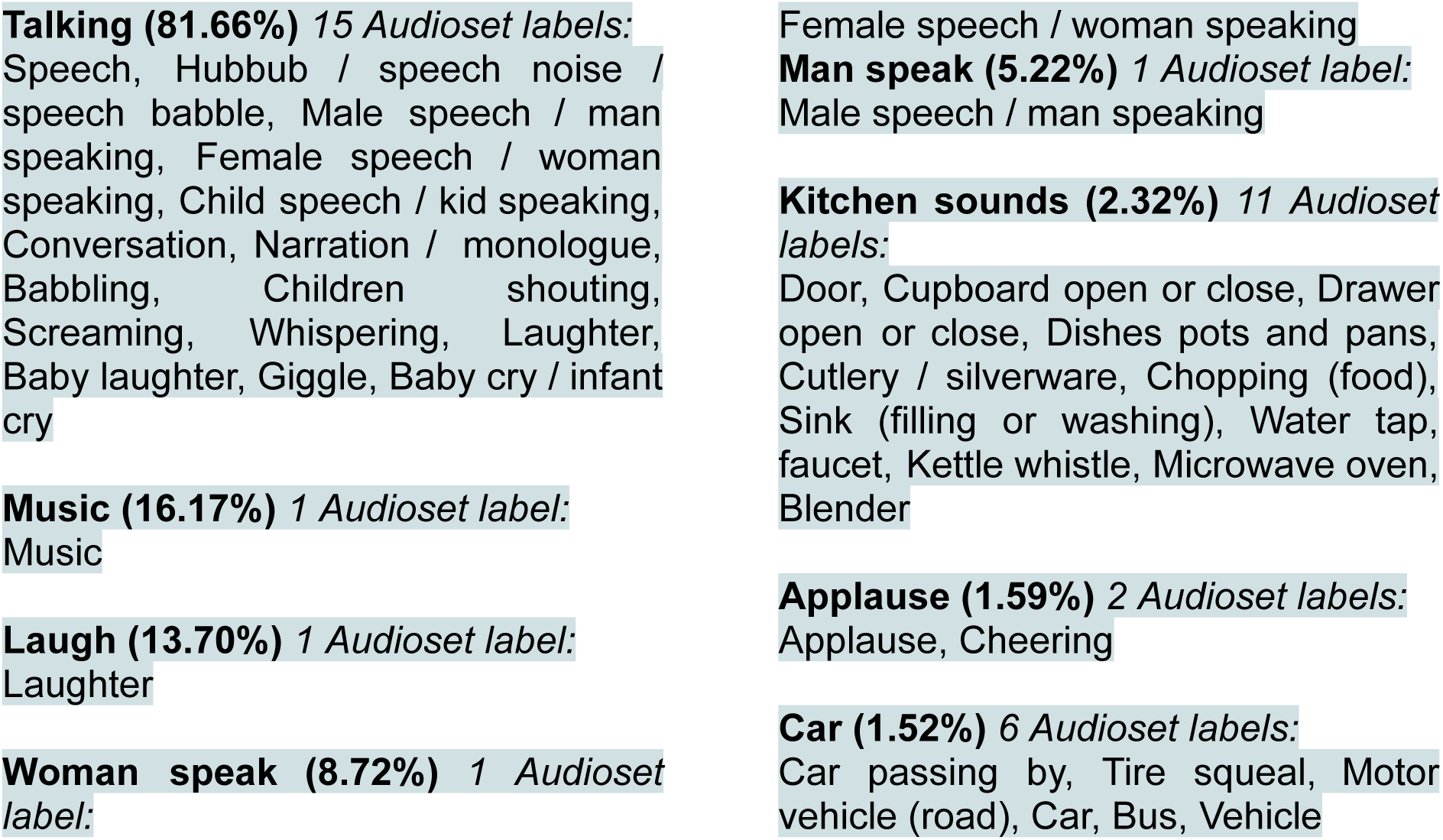
Categories used to label audio features in the *Friends* dataset. The categories are derived from AudioSet labels. Categories are in descending order, from Categorie most present in the audio of the dataset to least present (mean percentage through all 4 seasons, from top to bottom, left to right column).

### Appendix Results

#### Annex D Optimal hyperparameters substantially improve brain encoding with SoundNet

We optimised the hyperparameters for training our brain encoding model, which predicts individual brain activity using SoundNet internal representations. Through a systematic grid search, we explored the impact of several key hyperparameters, including learning rate, temporal window size, convolutional kernel size, weight decay, and early stopping parameters.

We tested 1296 configurations of hyperparameters on the fMRI data from sub-03. When ordering each configuration by the maximal r^2^ score obtained on the validation set for the Whole-brain baseline model, we observed an important gap between the first and last models: models with best r^2^ scores scored as high as 0.39, when models ranking 400 and below achieved a r^2^ score of 0.07 at best. A similar result has been found for the middle STG baseline model, with a maximal r^2^ score of 0.45, and a decrease of the score up to 0.05 for the worst performing models. This gap has also been observed when clustering the 100 best models. The agglomerative clustering of the r^2^ brain encoding maps using the centroid method with Euclidean distance computed 3 clusters of configurations for the Whole-brain baseline model: the maximal r^2^ score in the averaged predicted brain map of the best cluster, with 20 configurations, is 0.36, and the mean r2 score is 0.033, while for the second cluster, the scores drop respectfully to 0.29 and 0.013 (70 configurations), and to 5.10^-2^ and 8.10-4 for the last cluster (10 configurations).

Through these experiments, we identified an optimal set of hyperparameters that were then used for fine-tuning the models. Overall, our hyperparameter optimization process played a crucial role in enhancing the quality of brain encoding and selecting the most effective hyperparameter configuration for the subsequent analyses.

#### Annex E Conv4 fine-tuning leads to substantial improvements in brain encoding

**Annex Figure 1.**
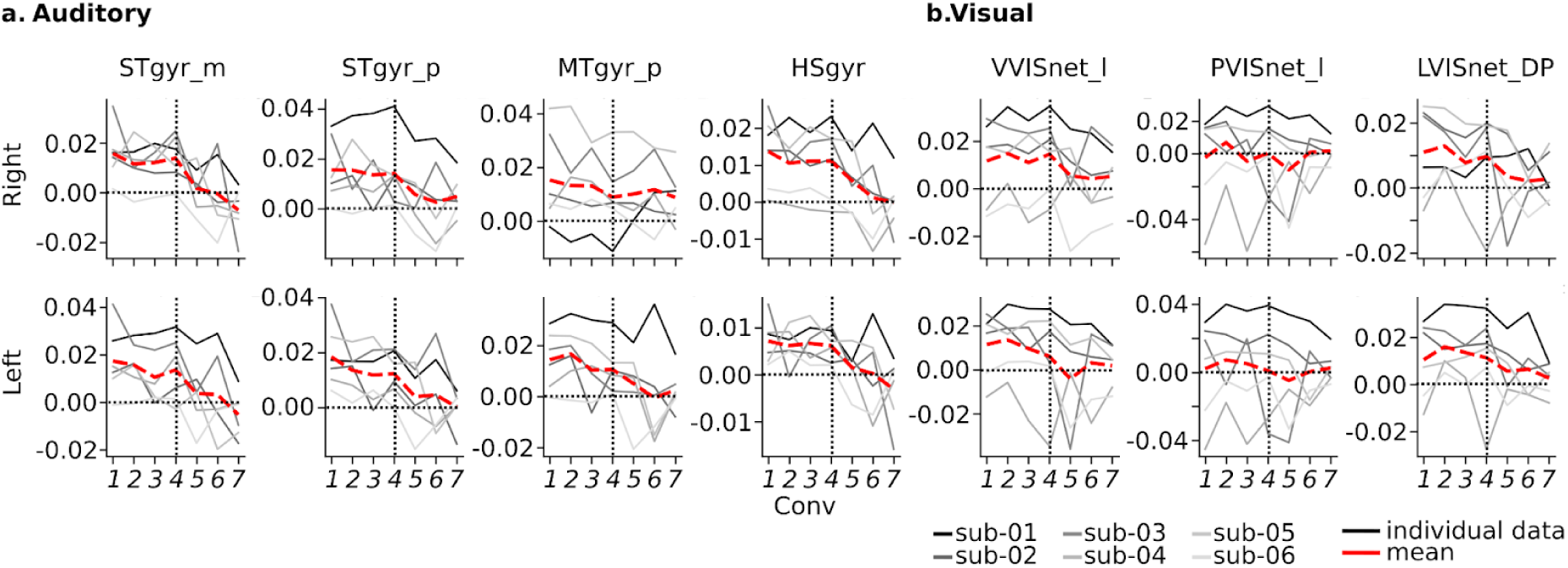
Impact of fine-tuning internal layers in SoundNet on individual brain encoding. Difference between the r² score (median across test runs) of the fine-tuned models (Conv1 to Conv7) minus baseline for each subject (sub-01 to sub-06) and the mean. Graphs are presented in the best predicted auditory (panel a) and visual (panel b) regions for the baseline model. STgyr_m: Superior Temporal gyrus middle; STgyr_p: Superior Temporal gyrus posterior; MTgyr_p: Middle Temporal gyrus posterior; HSgyr: Heschl’s gyrus; VVISnet_l: Ventral Visual network lateral; PVISnet_l: Posterior Visual network lateral; LVISnet_DP: Lateral Visual network dorsoposterior.

Our main goal was to see if fine-tuning an auditory model improved brain encoding performance. We adjusted SoundNet at different depths, using our training data (seasons 1 to 3), then tested the model using season 4. We calculated the median r² score for each of the 210 parcels of the MIST ROI atlas and averaged the performance across all subjects (Figure 3).

The best-predicted ROIs by the baseline model showed a consistent pattern (Figure 3). Adjusting models at Conv7, Conv6, or Conv5 did not improve brain encoding and even worsened it for the STG middle. However, models adjusted at Conv1 to Conv4 significantly improved over the baseline, with Conv4 showing the most substantial improvements. Adjusting at Conv1 to Conv3 only gave slight or no improvements over Conv4 for most ROIs, except the left lateral posterior visual network. We decided to focus on Conv4 models for further experiments based on these results.

#### Annex F Consistency of parcels best predicted between subjects

**Annex Figure 2.**
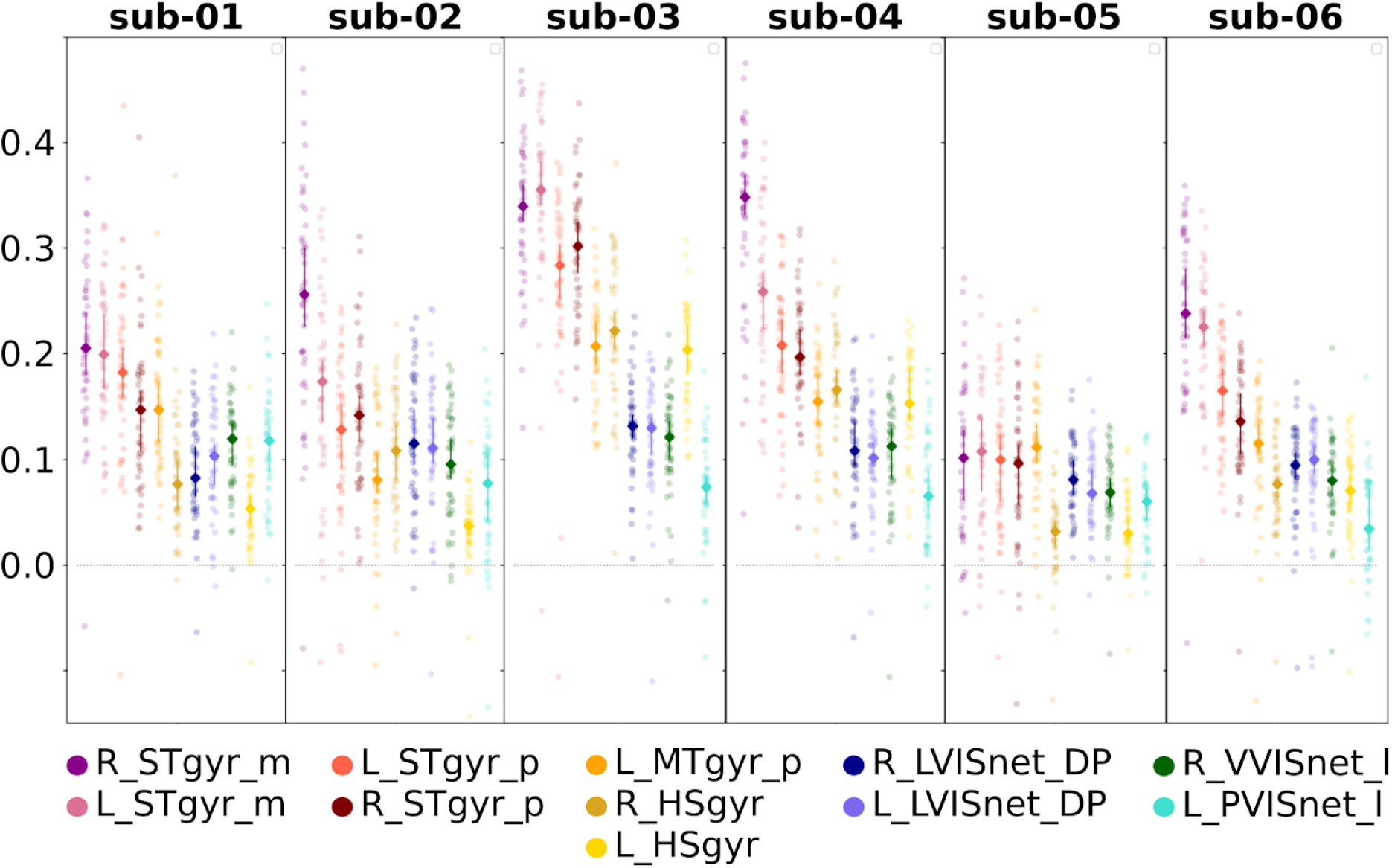
Distribution of r² scores across 48 runs (friends S04) for the eleven ROIs, with the highest average r² score across 6 subjects. The ROIs have been ordered by their median value across subjects and runs. The r² performance indicates the quality of prediction of fMRI time series in a single run (10 min. duration). R_STgyr_m: Right Superior Temporal gyrus middle; L_STgyr_m: Left Superior Temporal gyrus middle; L_STgyr_p: Left Superior Temporal gyrus posterior; R_STgyr_p: Right Superior Temporal gyrus posterior; L_MTgyr_p: Left Middle Temporal gyrus posterior; R_HSgyr: Right Heschl’s gyrus; L_HSgyr: Left Heschl’s gyrus; R_LVISnet_DP: Right Lateral Visual network dorsoposterior; L_LVISnet_DP: Left Lateral Visual network dorsoposterior; R_VVISnet_l: Right Ventral Visual network lateral; L_PVISnet_l: Left Posterior Visual network lateral.

#### Annex G Encoding brain activity at the voxel level without spatial smoothing

**Annex Figure 3.**
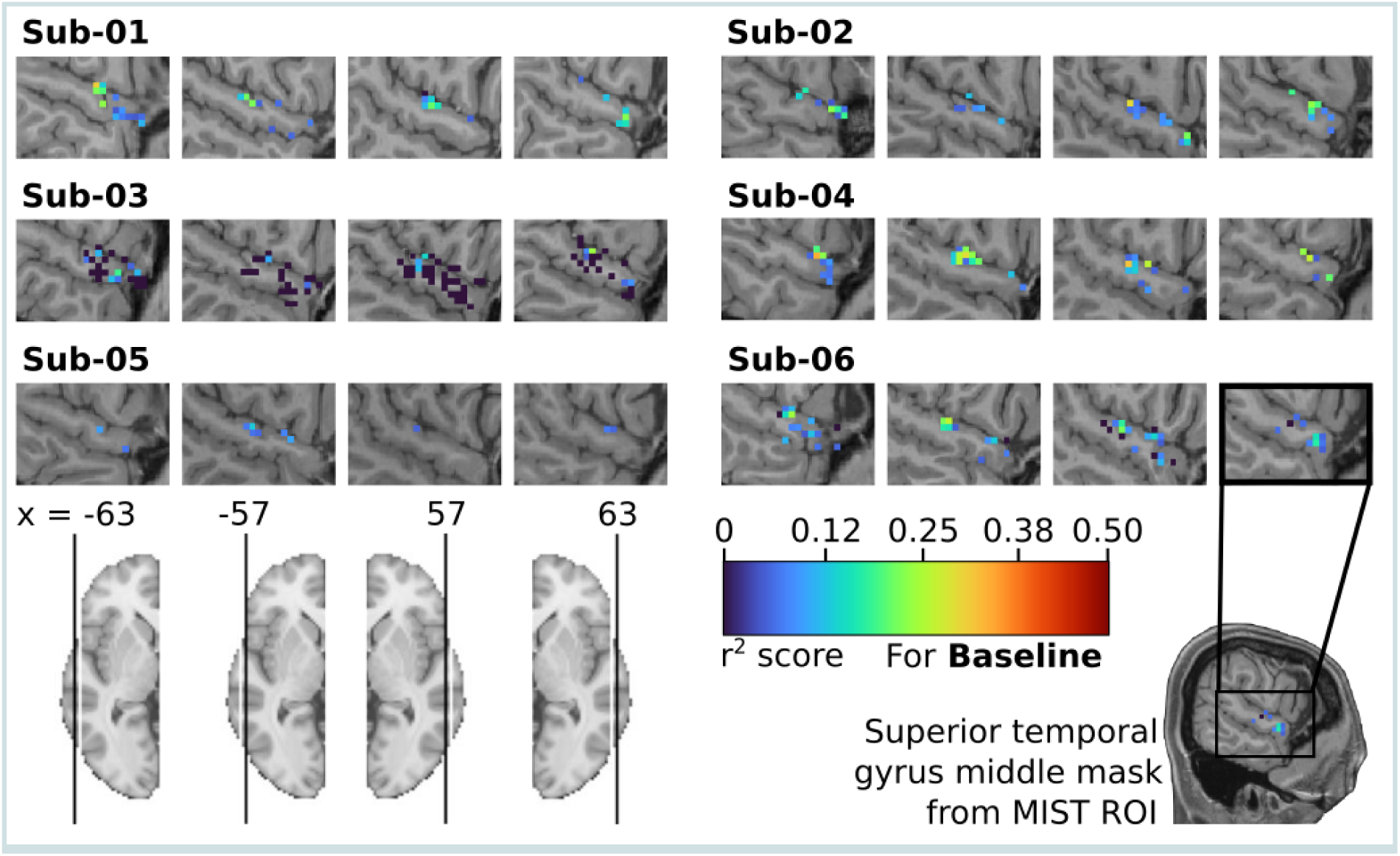
STG encoding using Soundnet with no fine-tuning and no spatial smoothing. Mapping of the r² scores from 556 voxels inside the cerebral region defined as the Middle STG by the parcellation MIST ROI, computed by the individual Baseline model. To have a better representation of the STG, 4 slices have been selected in each subject, 2 from the left hemisphere (-63 and -57) and 2 from the right hemisphere (63 and 57). Only voxels with r² values significantly higher than those of a null model initialised with random weights are shown (Wilcoxon test, FDR q < 0.05). Individual anatomical T1 have been used as background.

**Annex Figure 4.**
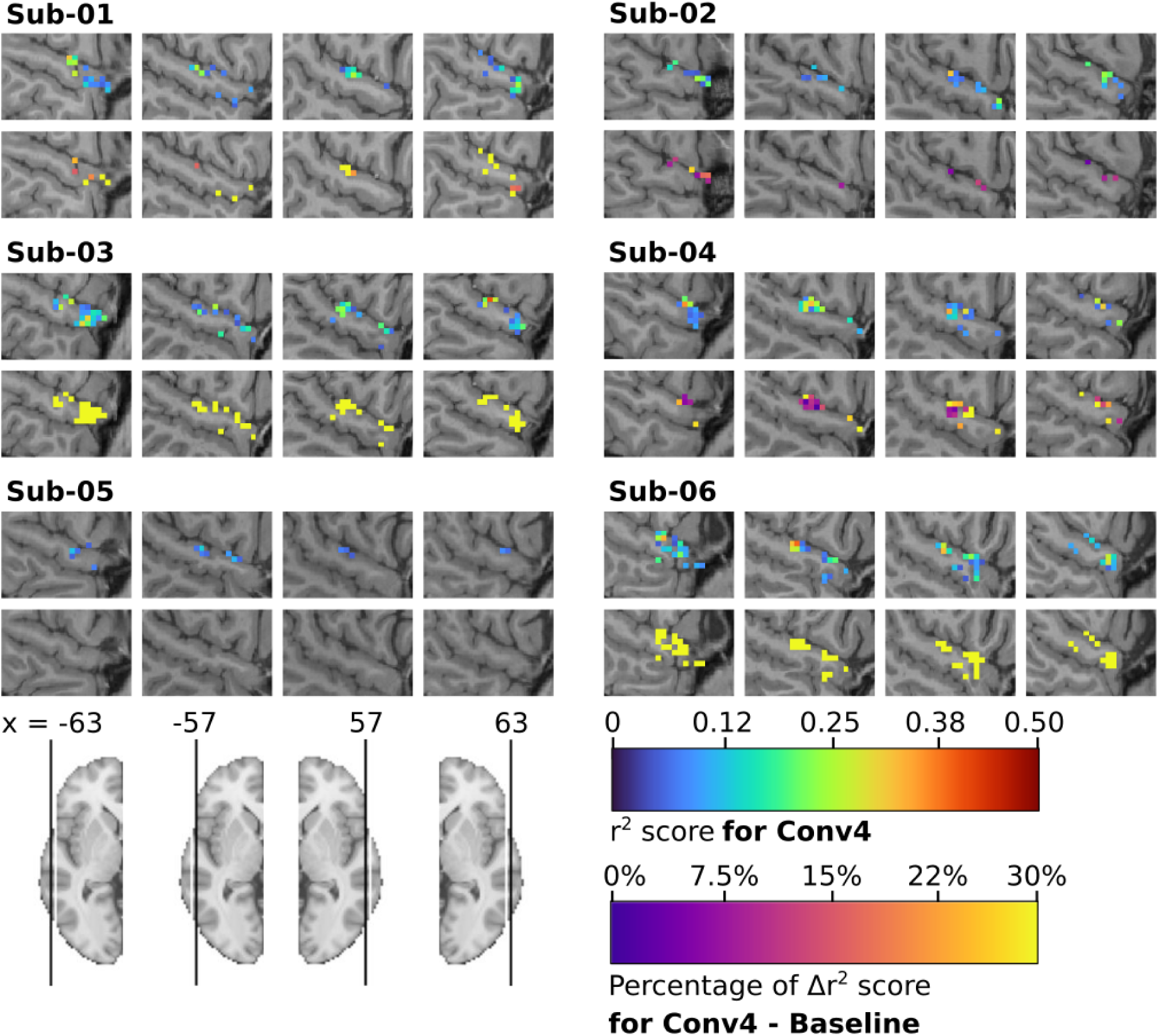
STG encoding using Brain-Aligned SoundNet and fMRI data with no spatial smoothing. For each subject, on the top : mapping of the r² scores from 556 voxels inside the cerebral region defined as the Middle STG by the parcellation MIST ROI, computed by the individual Conv4 model. Only voxels with r² values significantly higher than those of a null model initialised with random weights are shown (Wilcoxon test, FDR q < 0.05). For each subject, on the bottom : mapping of the difference of r² scores between the Conv4 model and the baseline model. Only voxels from the Conv4 model with r² values greater than +/- 0.05 and significantly greater or lesser than those of the baseline model are shown (Wilcoxon test, FDR q < 0.05). Individual anatomical T1 have been used as background.

#### Annex H Correlation between audio labels in Friends and change in prediction accuracy between individual baseline and brain-aligned models

**Annex Figure 5.**
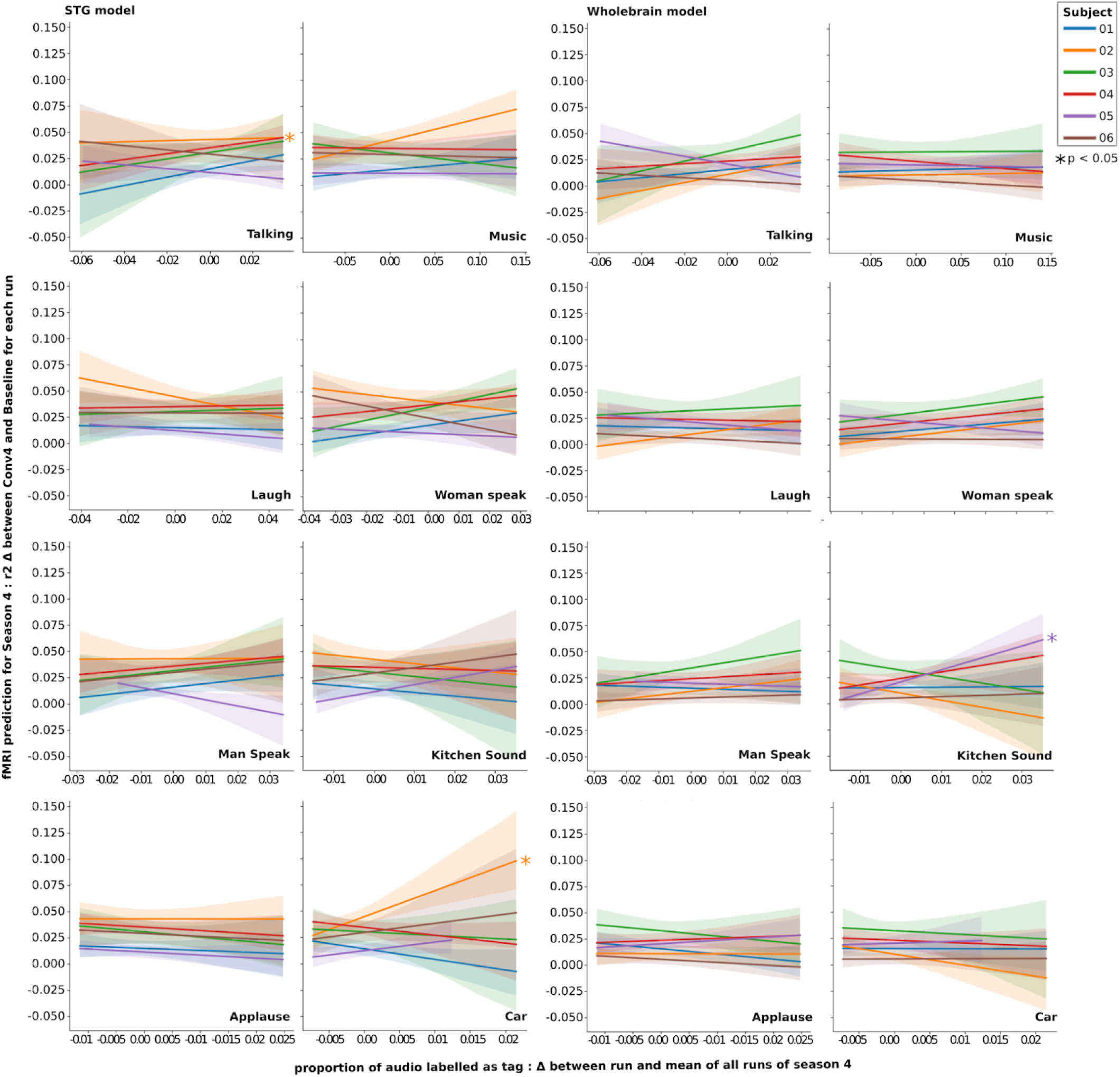
Linear regression of tagged audio proportion on prediction difference between brain-aligned and baseline model in season 4. Linear regression has been computed for each category, subject and model: We used the difference between the percentage of labelled audio of each half-episode and the mean percentage for all season 4 as the regressor, and the difference in max r² score between the baseline and brain-aligned models as the dependent variable. A multivariable regression (Ordinary Least Square) has also been computed for each subject and each model, using every category as regressors to explain the difference in max r² score. Significance (star) has been added to regressors with a significant (p < 0.05) in the OLS regression.

#### Annex I HEAR EVAL detailed score results

**Annex Figure 6.**
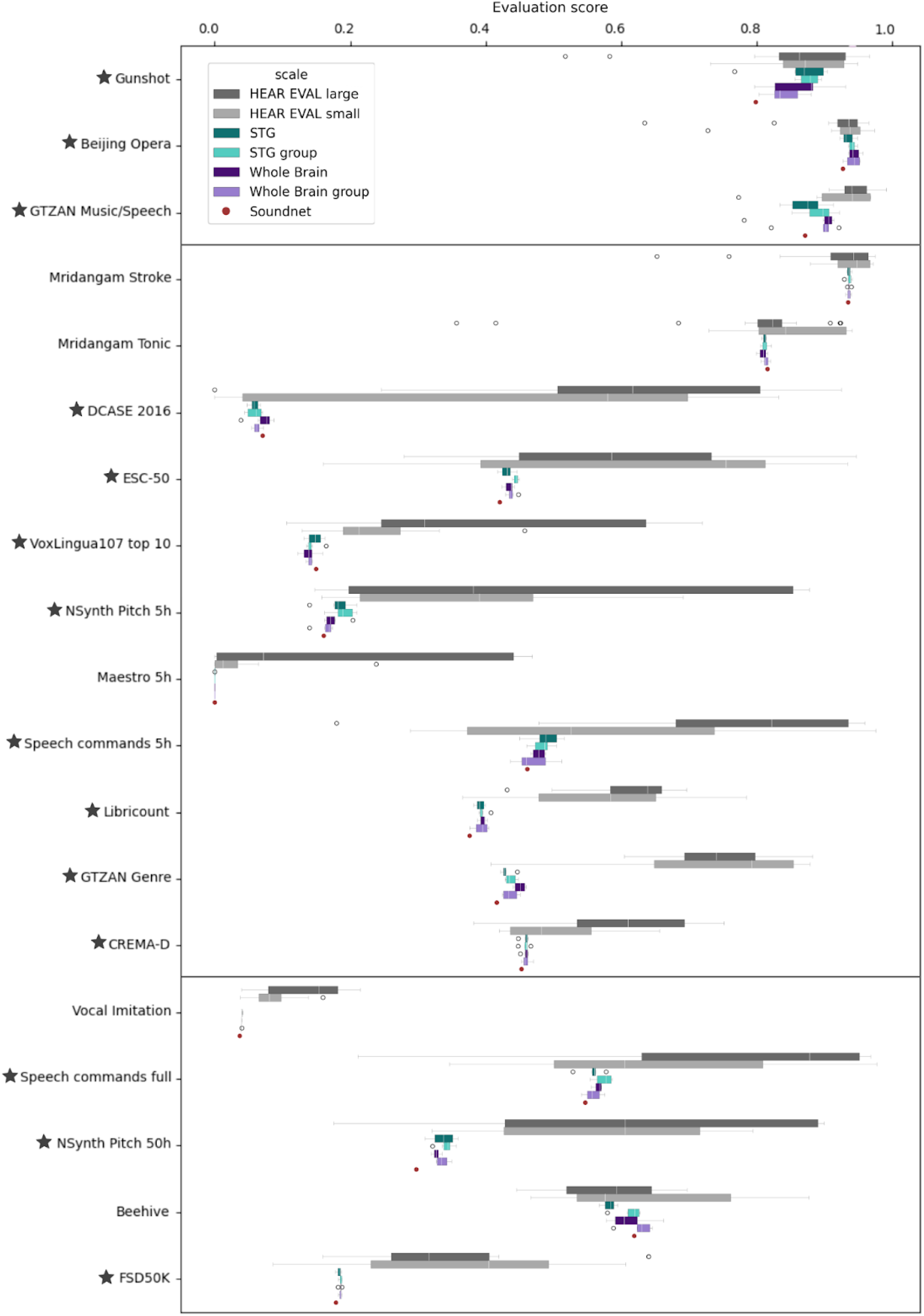
Distribution of all model’s scores. The models are grouped into the following categories based on their characteristics: *HEAR EVAL large* for models from the HEAR benchmark with more than 20 million parameters (up to 1339M), *HEAR EVAL small* for models from the HEAR benchmark with fewer than 12 million parameters, *STG* for models that are brain-aligned with fMRI data from the Superior Temporal Gyrus (STG) of a single subject, *Whole Brain* for models that are brain-aligned with a parcellation of the entire brain from a single subject, and respectively, *STG group* and *Whole Brain group* when using fMRI data from five subjects. *SoundNet* serves as the baseline, showing the performance of the pretrained SoundNet model, which is not brain-aligned. Tasks are ordered by size of the dataset used for training (estimation based on the HEAR paper) and separated in three categories: Small dataset (inferior to 1h, between 1h and 10h, more than 10h). Different metrics have been used depending on the task (Accuracy for classification, pitch accuracy, Onset only F-measure, aucroc, mAP). For each metric, a higher score relates to a better performance. For tasks annotated with a star, brain-aligned models performance is significantly different from SoundNet (Wilcoxon test, p < 0.05). Brain-aligned models significantly degrade network performance in two tasks only, DCASE 2016 and VoxLingua 107 top 10. In all other annotated tasks, performance has been improved.

## Footnotes

1 https://docs.cneuromod.ca/en/latest/index.html

2 https://github.com/SIMEXP/load_confounds

3 https://github.com/brain-bzh/cNeuromod_encoding_2020

4 https://github.com/smallflyingpig/SoundNet_Pytorch

5 https://hearbenchmark.com/hear-api.html

6 https://hearbenchmark.com/hear-tasks.html

7 https://hearbenchmark.com/hear-leaderboard.html

8 https://fmriprep.org/en/latest/workflows.html

9 https://docs.scipy.org/doc/scipy/reference/generated/scipy.cluster.hierarchy.linkage.html#scipy.cluster.hierarchy.linkage

