## Appendices - Methods & Results for "Alignment of auditory artificial networks with massive individual fMRI brain data leads to generalizable improvements in brain encoding and downstream tasks"

[1] https://fmriprep.org/en/latest/workflows.html

Annex B - Exploration of Hyperparameters space

| | | | | | *Early Stopping* | |
|---|---|---|---|---|---|---|
| *Sub* | Audio input length (tr) | Initial learning rate | Kernel size | Weight decay | Patience (epochs) | Delta |
| *03* | 1 | $1.10^{-2}$ | 1 | $1.10^{-2}$ | 10 | **0** |
| | 10 | $1.10^{-3}$ | 3 | **$1.10^{-3}$** | **15** | 0,1 |
| | 30 | **$1.10^{-4}$** | **5** | $1.10^{-4}$ | 20 | 0,5 |
| | **70** | | 9 | | | |
| *04* | 60 | $1.10^{-3}$ | 4 | | | |
| | 70 | $1.10^{-4}$ | 5 | | | |
| | **80** | **$1.10^{-5}$** | **6** | **$1.10^{-3}$** | **15** | **0** |
| *06* | 60 | $1.10^{-3}$ | 4 | | | |
| | **70** | $1.10^{-4}$ | 5 | | | |
| | 80 | **$1.10^{-5}$** | **6** | **$1.10^{-3}$** | **15** | **0** |
| *01* | 60 | $1.10^{-4}$ | 5 | | | |
| | 70 | **$1.10^{-5}$** | 6 | | | |
| | **80** | $1.10^{-6}$ | **7** | **$1.10^{-3}$** | **15** | **0** |
| *02* | 60 | $1.10^{-4}$ | 5 | | | |
| | 70 | **$1.10^{-5}$** | 6 | | | |
| | **80** | $1.10^{-6}$ | **7** | **$1.10^{-3}$** | **15** | **0** |
| *05* | **60** | $1.10^{-4}$ | 5 | | | |
| | 70 | $1.10^{-5}$ | 6 | | | |
| | 80 | **$1.10^{-6}$** | **7** | **$1.10^{-3}$** | **15** | **0** |

#### Annex C - Details of AudioSet categories used for audio annotation

**Talking (81.66%)** *15 Audioset labels:*
Speech, Hubbub / speech noise / speech babble, Male speech / man speaking, Female speech / woman speaking, Child speech / kid speaking, Conversation, Narration / monologue, Babbling, Children shouting, Screaming, Whispering, Laughter, Baby laughter, Giggle, Baby cry / infant cry

**Music (16.17%)** *1 Audioset label:*
Music

**Laugh (13.70%)** *1 Audioset label:*
Laughter

**Woman speak (8.72%)** *1 Audioset label:*
Female speech / woman speaking

**Man speak (5.22%)** *1 Audioset label:*
Male speech / man speaking

**Kitchen sounds (2.32%)** *11 Audioset labels:*
Door, Cupboard open or close, Drawer open or close, Dishes pots and pans, Cutlery / silverware, Chopping (food), Sink (filling or washing), Water tap, faucet, Kettle whistle, Microwave oven, Blender

**Applause (1.59%)** *2 Audioset labels:*
Applause, Cheering

**Car (1.52%)** *6 Audioset labels:*
Car passing by, Tire squeal, Motor vehicle (road), Car, Bus, Vehicle

**Annex Table 2. Categories used to label audio features in the *Friends* dataset.** The categories are derived from AudioSet labels. Categories are in descending order, from Categorie most present in the audio of the dataset to least present (mean percentage through all 4 seasons, from top to bottom, left to right column).

#### Appendix - Results

##### Annex D - Optimal hyperparameters substantially improve brain encoding with SoundNet

We optimised the hyperparameters for training our brain encoding model, which predicts individual brain activity using SoundNet internal representations. Through a systematic grid search, we explored the impact of several key hyperparameters, including learning rate, temporal window size, convolutional kernel size, weight decay, and early stopping parameters.

We tested 1296 configurations of hyperparameters on the fMRI data from sub-03. When ordering each configuration by the maximal $r^2$ score obtained on the validation set for the Whole-brain baseline model, we observed an important gap between the first and last models: models with best $r^2$ scores scored as high as 0.39, when models ranking 400 and below achieved a $r^2$ score of 0.07 at best. A similar result has been found for the middle STG baseline model, with a maximal $r^2$ score of 0.45, and a decrease of the score up to 0.05 for the worst performing models. This gap has also been observed when clustering the 100 best models. The agglomerative clustering of the $r^2$ brain encoding maps using the centroid method with Euclidean distance computed 3 clusters of configurations for the Whole-brain baseline model: the maximal $r^2$ score in the averaged predicted brain map of the best cluster, with 20 configurations, is 0.36, and the mean r2 score is 0.033, while for the second cluster, the scores drop respectfully to 0.29 and 0.013 (70 configurations), and to $5.10^{-2}$ and 8.10-4 for the last cluster (10 configurations).

Through these experiments, we identified an optimal set of hyperparameters that were then used for fine-tuning the models. Overall, our hyperparameter optimization process played a crucial role in enhancing the quality of brain encoding and selecting the most effective hyperparameter configuration for the subsequent analyses.

#### Annex E - Conv4 fine-tuning leads to substantial improvements in brain encoding

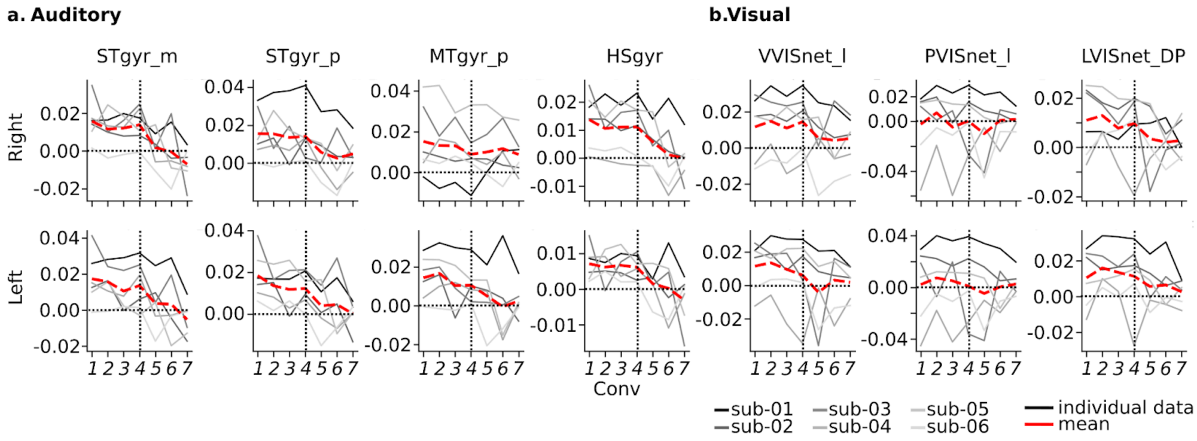


**Annex Figure 1. Impact of fine-tuning internal layers in SoundNet on individual brain encoding.** Difference between the r² score (median across test runs) of the fine-tuned models (Conv1 to Conv7) minus baseline for each subject (sub-01 to sub-06) and the mean. Graphs are presented in the best predicted auditory (panel a) and visual (panel b) regions for the baseline model. STgyr_m: Superior Temporal gyrus middle; STgyr_p: Superior Temporal gyrus posterior; MTgyr_p: Middle Temporal gyrus posterior; HSgyr: Heschl's gyrus; VVISnet_l: Ventral Visual network lateral; PVISnet_l: Posterior Visual network lateral; LVISnet_DP: Lateral Visual network dorsoposterior.

Our main goal was to see if fine-tuning an auditory model improved brain encoding performance. We adjusted SoundNet at different depths, using our training data (seasons 1 to 3), then tested the model using season 4. We calculated the median r² score for each of the 210 parcels of the MIST ROI atlas and averaged the performance across all subjects (Figure 3).

The best-predicted ROIs by the baseline model showed a consistent pattern (Figure 3). Adjusting models at Conv7, Conv6, or Conv5 did not improve brain encoding and even worsened it for the STG middle. However, models adjusted at Conv1 to Conv4 significantly improved over the baseline, with Conv4 showing the most substantial improvements. Adjusting at Conv1 to Conv3 only gave slight or no improvements over Conv4 for most ROIs, except the left lateral posterior visual network. We decided to focus on Conv4 models for further experiments based on these results.

#### Annex F - Consistency of parcels best predicted between subjects

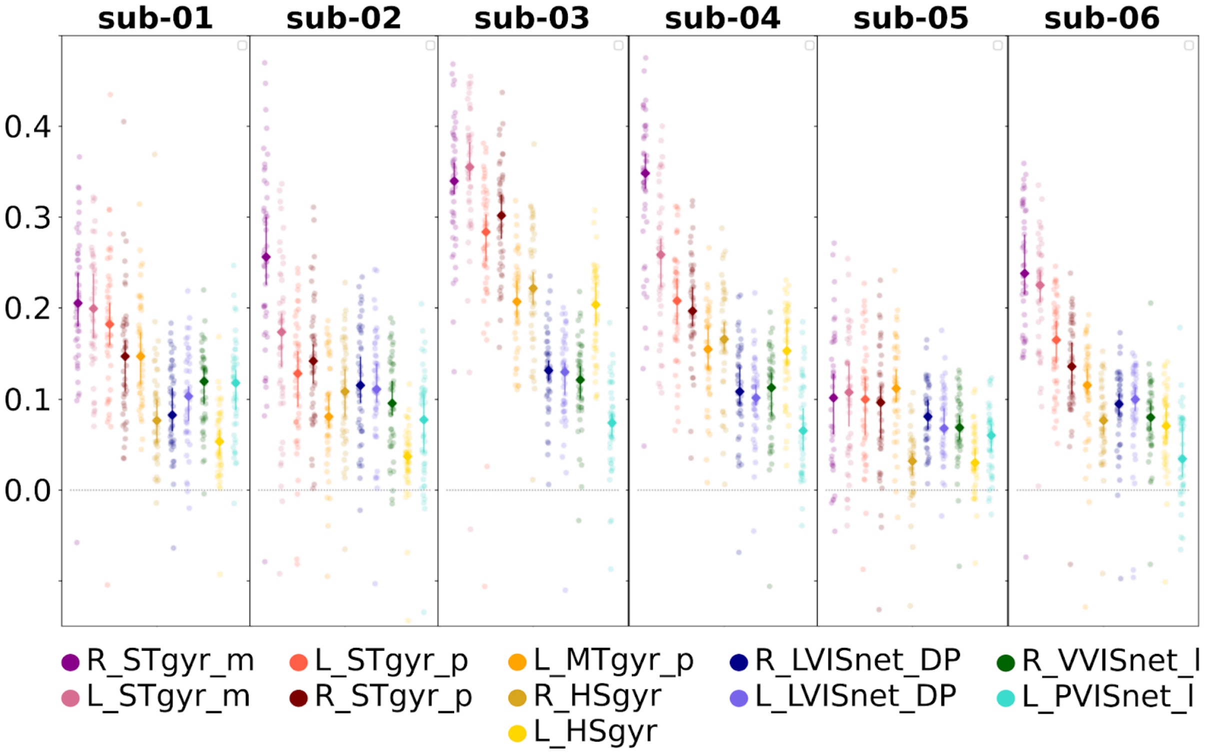


**Annex Figure 2.** Distribution of r² scores across 48 runs (friends S04) for the eleven ROIs, with the highest average r² score across 6 subjects. The ROIs have been ordered by their median value across subjects and runs. The r² performance indicates the quality of prediction of fMRI time series in a single run (10 min. duration). R_STgyr_m: Right Superior Temporal gyrus middle; L_STgyr_m: Left Superior Temporal gyrus middle; L_STgyr_p: Left Superior Temporal gyrus posterior; R_STgyr_p: Right Superior Temporal gyrus posterior; L_MTgyr_p: Left Middle Temporal gyrus posterior; R_HSgyr: Right Heschl's gyrus; L_HSgyr: Left Heschl's gyrus; R_LVISnet_DP: Right Lateral Visual network dorsoposterior; L_LVISnet_DP: Left Lateral Visual network dorsoposterior; R_VVISnet_l: Right Ventral Visual network lateral; L_PVISnet_l: Left Posterior Visual network lateral.

#### Annex G - Encoding brain activity at the voxel level without spatial smoothing

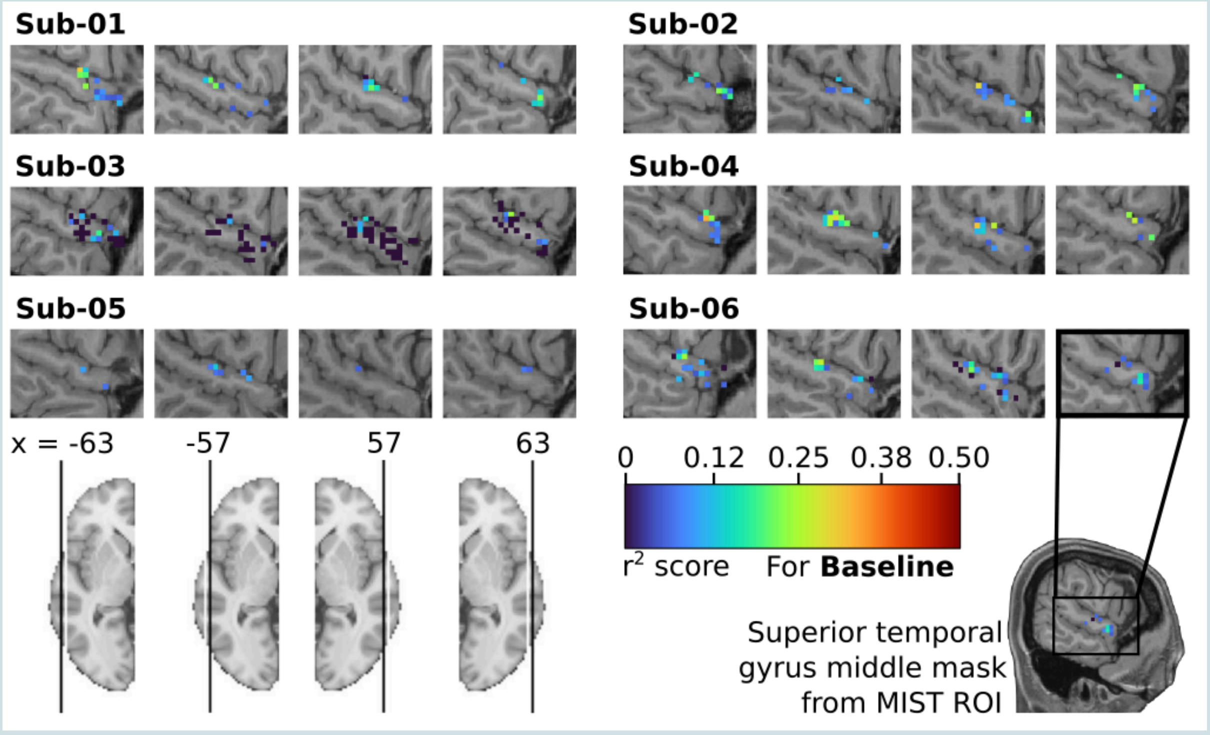


**Annex Figure 3. STG encoding using Soundnet with no fine-tuning and no spatial smoothing.** Mapping of the r² scores from 556 voxels inside the cerebral region defined as the Middle STG by the parcellation MIST ROI, computed by the individual Baseline model. To have a better representation of the STG, 4 slices have been selected in each subject, 2 from the left hemisphere (-63 and -57) and 2 from the right hemisphere (63 and 57). Only voxels with r² values significantly higher than those of a null model initialised with random weights are shown (Wilcoxon test, FDR $q < 0.05$). Individual anatomical T1 have been used as background.

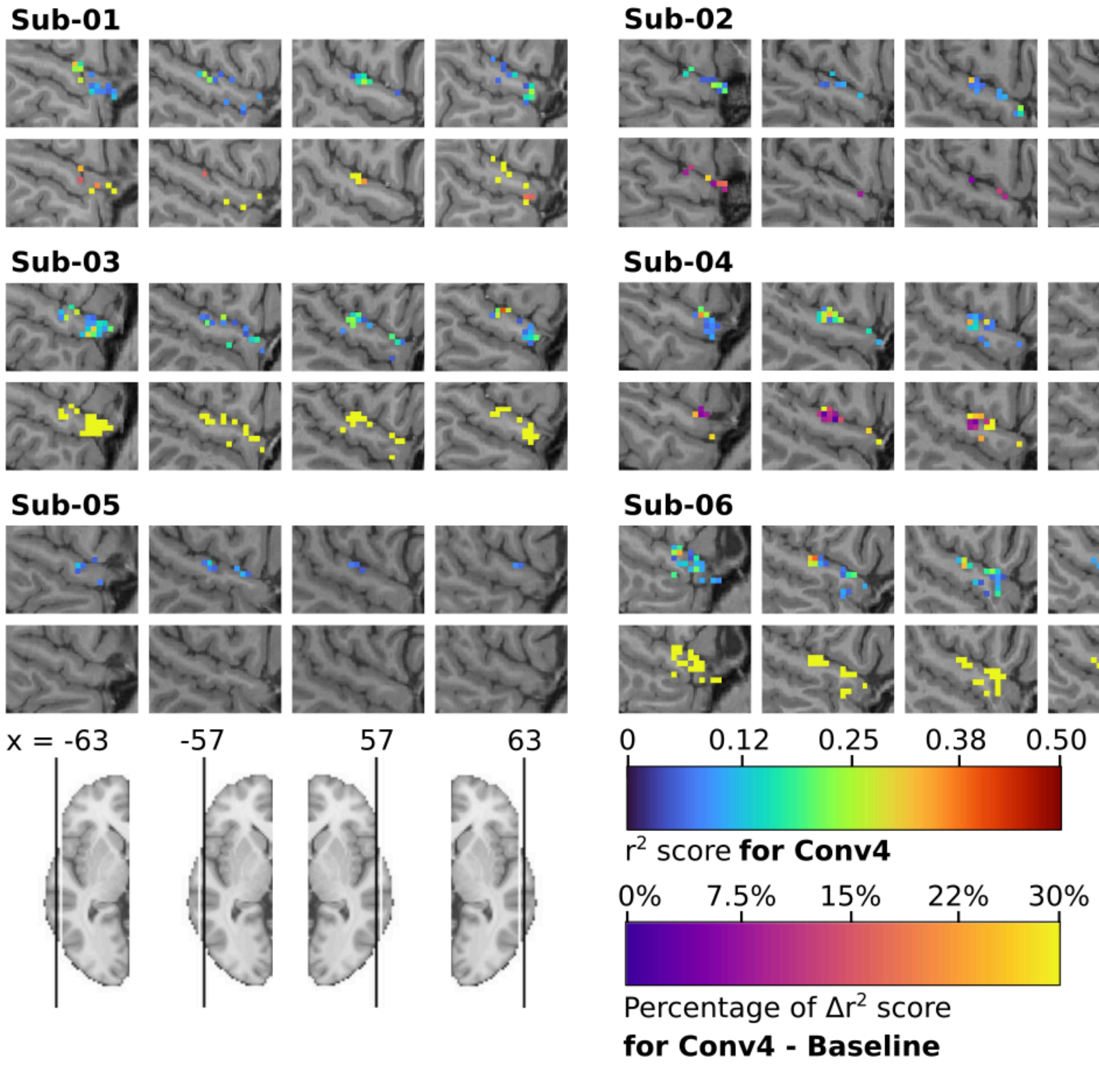


**Annex Figure 4. STG encoding using Brain-Aligned SoundNet and fMRI data with no spatial smoothing.** For each subject, on the top : mapping of the r² scores from 556 voxels inside the cerebral region defined as the Middle STG by the parcellation MIST ROI, computed by the individual Conv4 model. Only voxels with r² values significantly higher than those of a null model initialised with random weights are shown (Wilcoxon test, FDR q < 0.05). For each subject, on the bottom : mapping of the difference of r² scores between the Conv4 model and the baseline model. Only voxels from the Conv4 model with r² values greater than +/- 0.05 and significantly greater or lesser than those of the baseline model are shown (Wilcoxon test, FDR q < 0.05). Individual anatomical T1 have been used as background.

#### Annex H - Correlation between audio labels in Friends and change in prediction accuracy between individual baseline and brain-aligned models

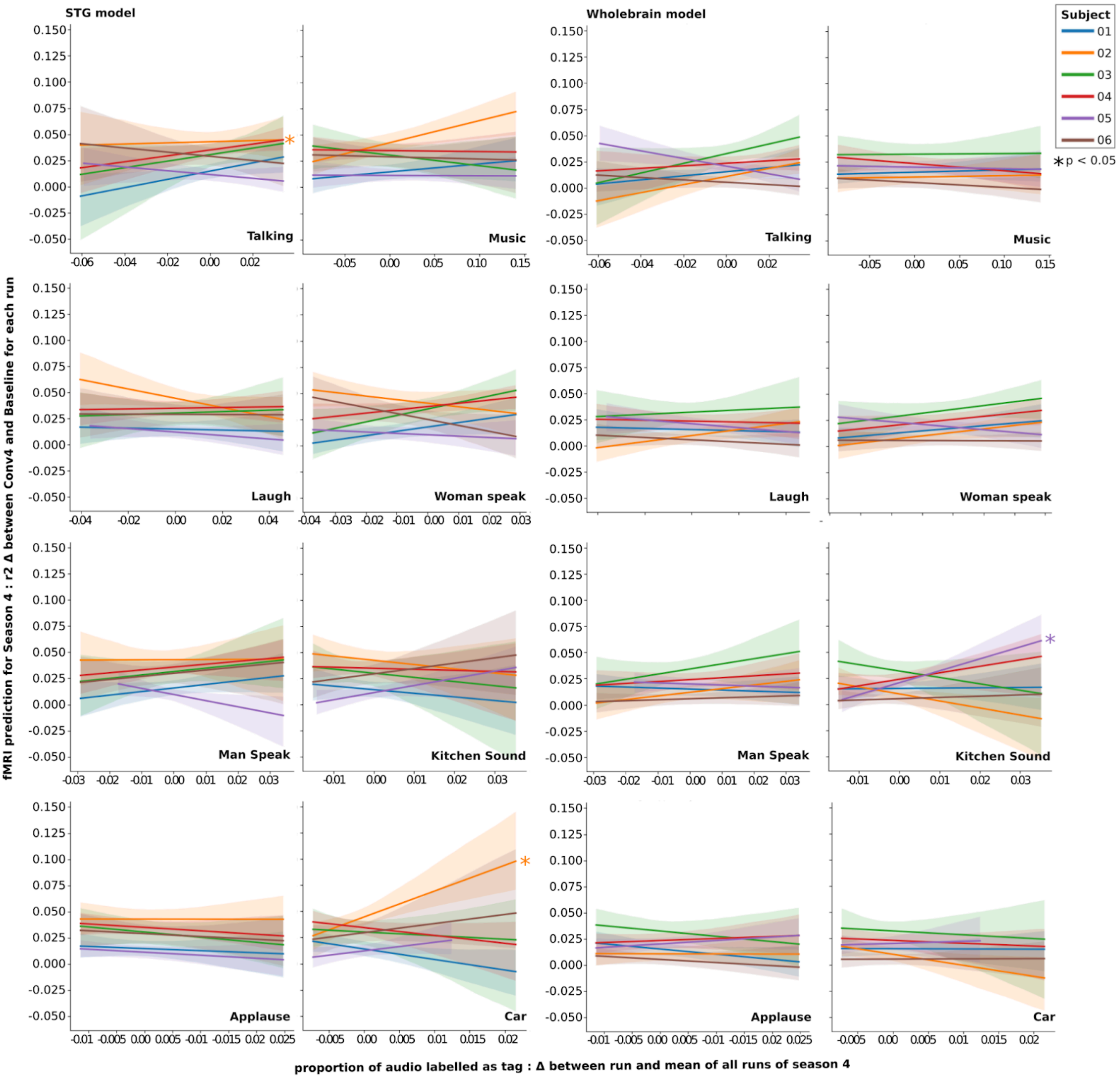


**Annex Figure 5. Linear regression of tagged audio proportion on prediction difference between brain-aligned and baseline model in season 4.** Linear regression has been computed for each category, subject and model: We used the difference between the percentage of labelled audio of each half-episode and the mean percentage for all season 4 as the regressor, and the difference in max r² score between the baseline and brain-aligned models as the dependent variable. A multivariable regression (Ordinary Least Square) has also been computed for each subject and each model, using every category as regressors to explain the difference in max r² score. Significance (star) has been added to regressors with a significant ($p < 0.05$) in the OLS regression.

### Annex I - HEAR EVAL detailed score results

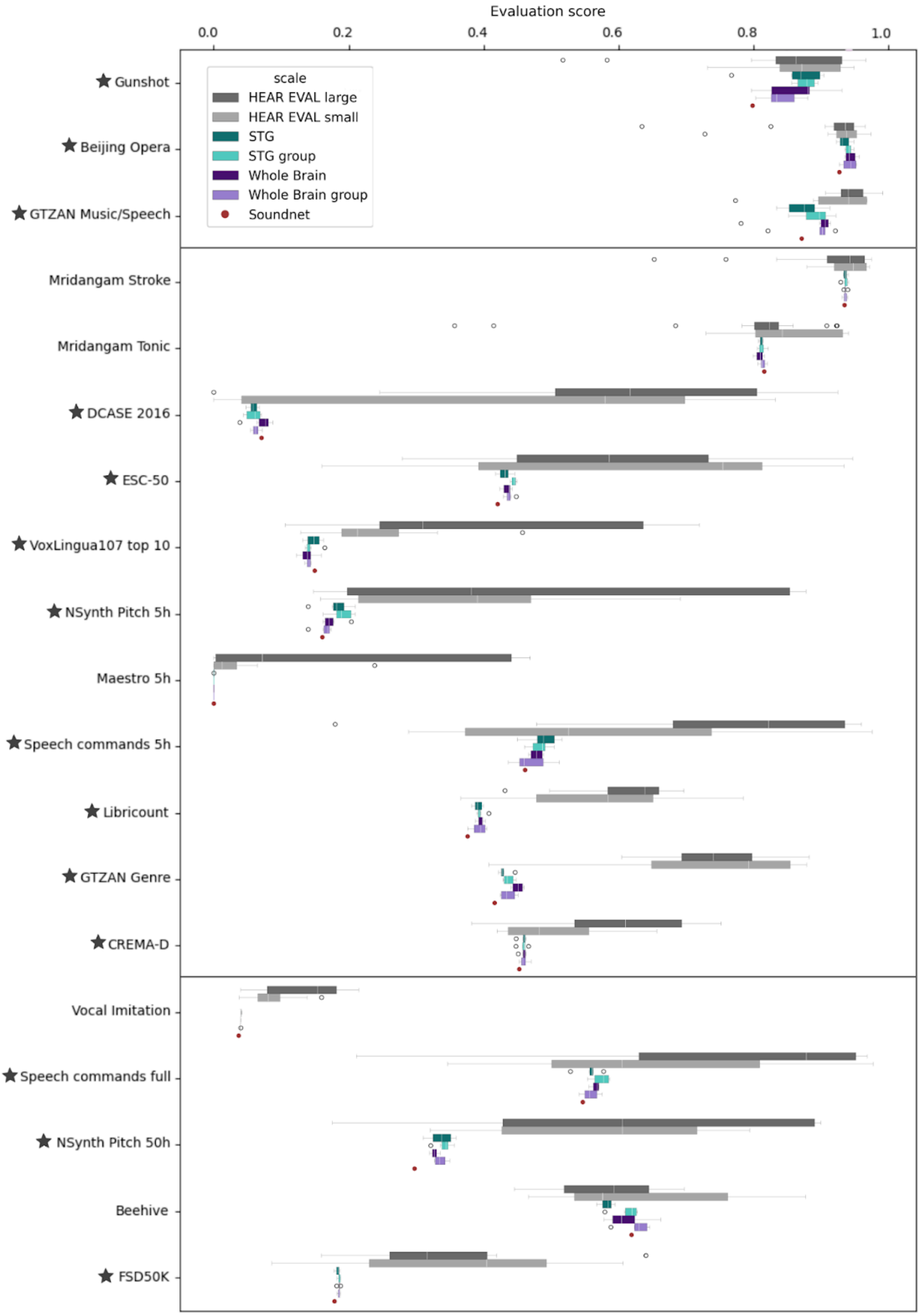

**Annex Figure 6. Distribution of all model's scores.** The models are grouped into the following categories based on their characteristics: *HEAR EVAL large* for models from the HEAR benchmark with more than 20 million parameters (up to 1339M), *HEAR EVAL small* for models from the HEAR benchmark with fewer than 12 million parameters, *STG* for models that are brain-aligned with fMRI data from the Superior Temporal Gyrus (STG) of a single subject, *Whole Brain* for models that are brain-aligned with a parcellation of the entire brain from a single subject, and respectively, *STG group* and *Whole Brain group* when using fMRI data from five subjects. *SoundNet* serves as the baseline, showing the performance of the pretrained SoundNet model, which is not brain-aligned. Tasks are ordered by size of the dataset used for training (estimation based on the HEAR paper) and separated in three categories: Small dataset (inferior to 1h, between 1h and 10h, more than 10h). Different metrics have been used depending on the task (Accuracy for classification, pitch accuracy, Onset only F-measure, aucroc, mAP). For each metric, a higher score relates to a better performance. For tasks annotated with a star, brain-aligned models performance is significantly different from SoundNet (Wilcoxon test, $p < 0.05$). Brain-aligned models significantly degrade network performance in two tasks only, DCASE 2016 and VoxLingua 107 top 10. In all other annotated tasks, performance has been improved.
